# Notch signaling maintains a progenitor-like subclass of hepatocellular carcinoma

**DOI:** 10.1101/2024.12.13.628320

**Authors:** Kerstin Seidel, Robert Piskol, Thi Thu Thao Nguyen, Amy Shelton, Charisa Cottonham, Cecile C. de la Cruz, Joseph Castillo, Jesse Garcia, Udi Segal, Mark Merchant, Yeqing Angela Yang, Jasmine Chen, Musa Ahmed, Alexis Scherl, Rajesh Vij, Lluc Mosteiro, Yan Wu, Zora Modrusan, Ciara Metcalfe, Chris Siebel

## Abstract

Hepatocellular carcinomas (HCCs) constitute one of the few cancer indications for which mortality rates continue to rise. While Notch signaling dictates a key progenitor lineage choice during development, its role in HCC has remained controversial. Using therapeutic antibodies targeting Notch ligands and receptors to screen over 40 patient-derived xenograft models, we here identify progenitor-like HCCs that crucially depend on a tumor-intrinsic JAG1-NOTCH2 signal. Inhibiting this signal induces tumor regressions by triggering progenitor-to-hepatocyte differentiation, the same cell fate-switch that Notch controls during development. Transcriptomic analysis places the responsive tumors within the well-characterized progenitor subclass, a poor prognostic group of highly proliferative tumors, providing a diagnostic method to enrich for Notch-dependent HCCs. Furthermore, single-cell RNA sequencing uncovers a heterogeneous population of tumor cells and reveals how Notch inhibition shifts cells from a mixed cholangiocyte-hepatocyte lineage to one resembling mature hepatocytes. Analyzing the underlying transcriptional programs brings molecular detail to this process by showing that Notch inhibition de-represses expression of CEBPA, which enables the activity of HNF4α, a hepatocyte lineage factor that is otherwise quiescent. We thus describe a compelling and targetable dependency in a poor-prognosis class of HCCs.

## Introduction

The liver employs two main types of epithelial cells---hepatocytes, the predominant parenchymal cell type, and cholangiocytes, which line the bile ducts---to provide key metabolic functions, including the synthesis and secretion of bile acids and detoxification (Stanger, 2015). Early embryonic development (E8-E9.5) in mammals integrates signals from multiple signaling pathways, notably Wnt, FGF and BMP, to initiate and drive hepatic development, including the generation of embryonic bi-potent progenitor cells called hepatoblasts (Si-Tayeb et al., 2010). Active Notch signals direct hepatoblasts to differentiate to the cholangiocyte fate whereas an absence of such signals enables hepatocyte differentiation (Adams & Jafar-Nejad, 2019). Downstream of Notch and these other signaling pathways, transcription factor networks control cell fate, proliferation and differentiation; in particular, HNF4A functions centrally in hepatoblast specification, with loss or gain of CEBPA determining cholangiocyte versus hepatocyte fate, respectively (Tachmatzidi et al., 2021).

Liver cancers rank among the world’s most commonly diagnosed cancers and are the third leading cause of cancer mortalities (Sung et al., 2021). Hepatocellular carcinomas (HCCs), which constitute the majority of liver cancers, primarily arise in the context of cirrhosis, which is commonly caused by Hepatitis B or C virus (HBV, HCV) infections, toxin exposure, alcoholism, or non-alcoholic steatohepatitis (NASH) (Llovet et al., 2022). HCC is commonly diagnosed at advanced stages. Curative resection or ablation is possible only in early stage tumors and recurrence rates are high (Llovet et al., 2022). Systemic therapies now include tyrosine kinase inhibitors (da Fonseca et al., 2020) as well as the combination of the therapeutic antibodies atezolizumab and bevacizumab to block immune checkpoints and angiogenesis, respectively (Finn et al., 2020). While HCC management has improved patient outcomes over the past decade, there remains a strong medical need to develop improved adjuvant therapies and increase the efficacy and duration of systemic therapies (Llovet et al., 2022).

The term “HCC” serves as an umbrella classifier for a heterogenous cancer type composed of multiple subtypes that reflect differences in etiology, cell of origin, and driver signaling pathways and mutations. Histological classifications broadly categorize tumors as poorly- or well-differentiated and further sub-group tumors based on content of cholangiocyte or hepatocyte features, including mixed-lineage and progenitor tumors (Calderaro et al., 2019). These features have also been correlated with different types of molecular classifiers. The most frequent genetic mutations highlight dysregulated cell cycle control, telomerase activation, Wnt signaling, epigenetic regulation and oncogenic tyrosine kinase (as evidenced in part by *FGF19*, *VEGFA*, and *MET* chromosomal amplifications) and MAPK signaling (Llovet et al., 2022). Transcriptome meta-analyses of global patient populations have helped classify HCCs into proliferative and non-proliferative groups, which largely correspond to the poorly- and well-differentiated classes, respectively. Within the proliferative population, several groups have molecularly identified a progenitor subclass (Ally et al., 2017; Boyault et al., 2007; Hoshida et al., 2009), characterized by the expression of stem/progenitor cell markers (Hoshida et al., 2009) and an aggressive, poor-prognosis phenotype (Llovet et al., 2022).

Notch signals, conserved throughout metazoans, vitally contribute to specifying the fate and function of neighboring cells, including progenitors during liver development (Siebel & Lendahl, 2017). In mammals, binding of cell surface ligands of the Jagged (Jag) or Delta-like (Dll) families conformationally alters the negative-regulatory region (NRR) of one of four Notch receptors (NOTCH1-4) (Gordon et al., 2015) to ultimately trigger activation through a gamma-secretase-catalyzed proteolytic cleavage (De Strooper et al., 1999). The released NOTCH intracellular domain (NICD) translocates into the nucleus, assembles into a transcription factor complex that includes the DNA-binding protein RBPJ, and activates the downstream transcriptional program. Given that Notch signaling has been linked to numerous human diseases, including cancer, inflammation and metabolic disorders (Allen & Maillard, 2021; Aster et al., 2017; Xu & Wang, 2021), researchers have developed numerous Notch-blocking therapeutics, including gamma-secretase inhibitors and monoclonal antibodies that selectively target individual Notch ligands and receptors (Allen & Maillard, 2021).

Mutations in Notch receptors that activate or prolong Notch signaling point to an oncogenic role in multiple cancers, including T cell acute lymphoblastic leukemia (T-ALL), triple-negative breast cancer and adenoid cystic carcinoma (Aster et al., 2017). However, such mutations have not been broadly described in liver cancers, and conflicting reports obscure a clear understanding of Notch signaling in HCC. Whereas some reports suggest that Notch signals suppress HCC development (Viatour et al., 2011), other studies conclude that hepatocyte Notch signaling can drive fibrosis in NASH (Zhu et al., 2018), oncogenesis in HCC (Dill et al., 2013) and expression of a conserved gene signature in HCC patient populations (Villanueva et al., 2012). Furthermore, achieving the therapeutically vital goal of defining which Notch ligands and receptors function in HCC requires moving past gain-of-function methods to approaches that precisely inhibit individual ligands and receptors in physiologically relevant preclinical models.

By conducting a preclinical trial of a JAG1-selective blocking antibody in a large panel of patient-derived xenograft models, we now report the discovery of HCCs that exquisitely depend on a JAG1-NOTCH2 signaling axis within tumor cells. Molecular characterization shows that these tumors display hallmarks of embryonic liver progenitors in the aggressive, proliferative HCC subclass. Strikingly, selective inhibition of JAG1 or NOTCH2 leads to profound tumor regressions. Single-cell sequencing analyses of these tumors and their responses following treatment reveal a mechanism in which Notch inhibition induces cycle arrest and promotes hepatocyte differentiation, through upregulation of CEBPA expression and activation of existing HNF4A, mimicking normal developmental programs. Our results thus identify a Notch-dependent subset of HCCs within the progenitor subclass, outline the relevant transcriptional and differentiation mechanisms, and highlight JAG1 and NOTCH2 as actionable targets in this poor-prognosis group.

## Results

### Discovery of a Notch-dependent HCC subclass

Previous work from our group implicated JAG1 and NOTCH2 as a functionally relevant Notch ligand-receptor pair that supported growth of HCC and cholangiocarcinoma (CC) tumor types (Huntzicker et al., 2015a). However, this study largely relied on the *in vivo* transfection of mouse liver cells with constitutively active N-Ras and Akt. To test the hypothesis that JAG1 and NOTCH2 were drivers of HCC in a pre-clinical system using patient samples, we moved to patient-derived xenografts (PDXs) involving the direct transfer of tumor samples from patient to an immunocompromised mouse (Siolas & Hannon, 2013). Relying on transcriptome data that showed expression of *JAG1* and *NOTCH2* as well as evidence of pathway activity, such as expression of *HES1*, we chose two models for initial tests of sensitivity to aJ1.b70, a potent and selective therapeutic antibody that inhibits JAG1 (Lafkas et al., 2015). While JAG1 blockade had no effect on the growth of one PDX model, aJ1.b70 treatment potently inhibited growth of the second model, LIV78, in a dose-dependent manner (Fig. 1A). This striking response---regression of an established solid tumor, induced by a single-agent targeting Notch signaling---was unprecedented in our experience of manipulating Notch signaling in solid cancers.

**Figure 1.**
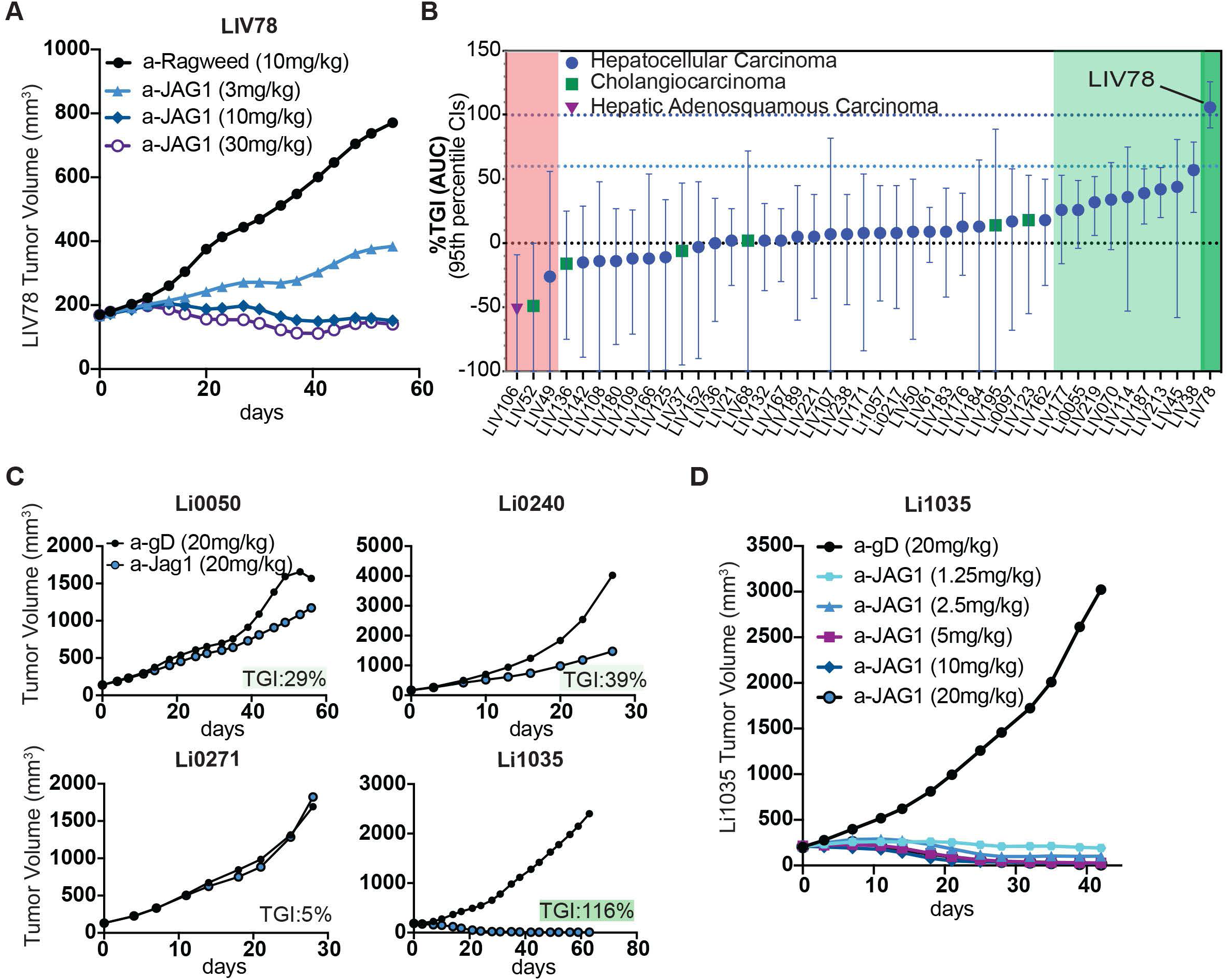
Discovery of HCC PDX tumors that are highly sensitive to JAG1 inhibition. **A)** *In vivo* efficacy of aJ1.b70 treatment in LIV78 HCC PDX model. Data is shown for multiple doses. **B)** Analysis of tumor growth inhibition in an unbiased panel of liver tumor PDX models following aJ1.b70 treatment. Dark green panel, complete response; light green panel, partial response, white panel, no response; red panel, increased tumor growth. **C)** *In vivo* efficacy of aJ1.b70 treatment in models selected based on shared features with LIV78. Extent of growth response is highlighted as in B) within each subpanel. **D)** *In vivo* efficacy of aJ1.b70 treatment HCC PDX model Li-1035. Data is shown for multiple doses.

Given that we discovered this Notch-dependent tumor in one of two PDX models using simple screening criteria and that the majority of PDX models and HCC tumors express JAG1 and NOTCH2 (Huntzicker et al., 2015a), TCGA Research Network: https://www.cancer.gov/tcga), we initiated a pre-clinical trial to assess the frequency of response to Notch inhibition. We tested aJ1.b70 in a panel of 42 additional PDX liver cancer models (comprising 35 HCC models plus seven CCs and other liver tumor subtypes), chosen without bias or prescreening, including for Notch pathway status. We used five mice per control and anti-JAG1 groups and dosed for approximately three weeks, given the ambitious scope of the study. Examining the percentage of aJ1.b70 tumor growth inhibition (%TGI) revealed that Jag1 blockade had little to no effect on growth of 30 tumors, although %TGI varied significantly between individual tumors, likely reflecting the small group sizes and short growth period (Fig. 1B, white panel). Three models, including a CC and an adenosquamous carcinoma, trended towards enhanced growth following treatment (Fig. 1B, red area). We classified 9 tumors as partial responders, with %TGI ranging from approximately 26 to 57 (Fig. 1B, light green area). Thus, this unbiased *in vivo* screen revealed only a modest JAG1-dependence (21%) within this cohort of PDX. LIV78 (dark green area), however, strikingly stood out as showing treatment-induced regressions.

We expanded our screen, searching for additional “super-responder” models that shared features of LIV78. In particular, LIV78 not only expressed high levels of *JAG1* and *NOTCH2*, but also expressed multiple markers of embryonic liver progenitor cells. Using this approach, we identified four additional models and tested them for sensitivity to aJ1.b70 *in vivo*. Three models responded moderately or not at all (Fig. 1C). However, one model, Li1035, responded dramatically (Fig. 1C), showing tumor regressions in a dose-dependent manner (Fig. 1D) and revealing a JAG1 dependence similarly strong as that observed in LIV78.

To identify the Notch signaling components expressed in both the human cancer cells and mouse stromal cells that typically comprise PDX tumors, we performed bulk RNA sequencing. Tumors from LIV78 and Li1035 were composed of approximately 95% human cancer cells and 5% mouse stromal cells, based on the percentages of human versus mouse sequencing reads (Supplemental Fig. 1A). Among the four canonical ligands, *JAG1* was the only one expressed by the human cancer cells (Fig. 2A), consistent with growth inhibition following JAG1 blockade. In contrast, *Dll4* was the only ligand significantly expressed in mouse cells, likely reflecting the well-characterized DLL4 expression in endothelial cells (Mack & Luisa Iruela-Arispe, 2018) (Fig. 2A). Both LIV78 and Li1035 cancer cells expressed human *NOTCH1* and *NOTCH2* whereas the mouse stromal cells expressed *Notch1*, *Notch3* and *Notch4* (Fig. 2A). To determine whether NOTCH1, NOTCH2 or both were the relevant receptors mediating Notch dependency, we treated LIV78 or Li1035 PDX models with blocking antibodies that selectively and potently inhibit each receptor (Wu et al., 2010). Similar to treatment with aJ1.b70, treatment with a NOTCH2 blocking antibody (aNRR2) induced regressions of LIV78 and Li1035 tumors (Figs. 2B and C) whereas a NOTCH1 blocking antibody (aNRR1) showed only a modest and delayed effect (Fig. 2C). Immunohistochemical staining with an antibody targeting an epitope in the NOTCH2 intracellular domain (NICD2, cleaved and uncleaved forms) revealed clear NOTCH2 staining in the majority of tumor cells in both models, including membranous/diffuse (inactive) and nuclear (active) staining. Active (nuclear) NOTCH2 signaling was observed in approximately 30-50% of tumor cells. (Fig. 2D.i and Supplemental Fig. 1B.i), while nuclear NICD2 staining was not observed in non-regressing models (Fig. 1C, Supplemental Fig. 1C). Most importantly, tumors harvested after treatment with aJ1.b70 or aNRR2 no longer showed nuclear NICD2 staining, consistent with effective blockade of the Notch2 signal (Fig. 2D.ii-iii and Supplemental Fig. 1B.ii-iii). In contrast, Notch1 signaling was detected only in what morphologically appeared to be endothelial cells (Fig. 2D.v-viii and Supplemental Fig. 1B.v-viii); and treatment with aNRR1 inhibited the endothelial Notch1 signal (Fig. 2D.viii and Supplemental Fig. 1B.viii) without affecting the NICD2 nuclear signal in tumor cells (Fig. 2D.iv and Supplemental Fig. 1B.iv).

**Figure 2.**
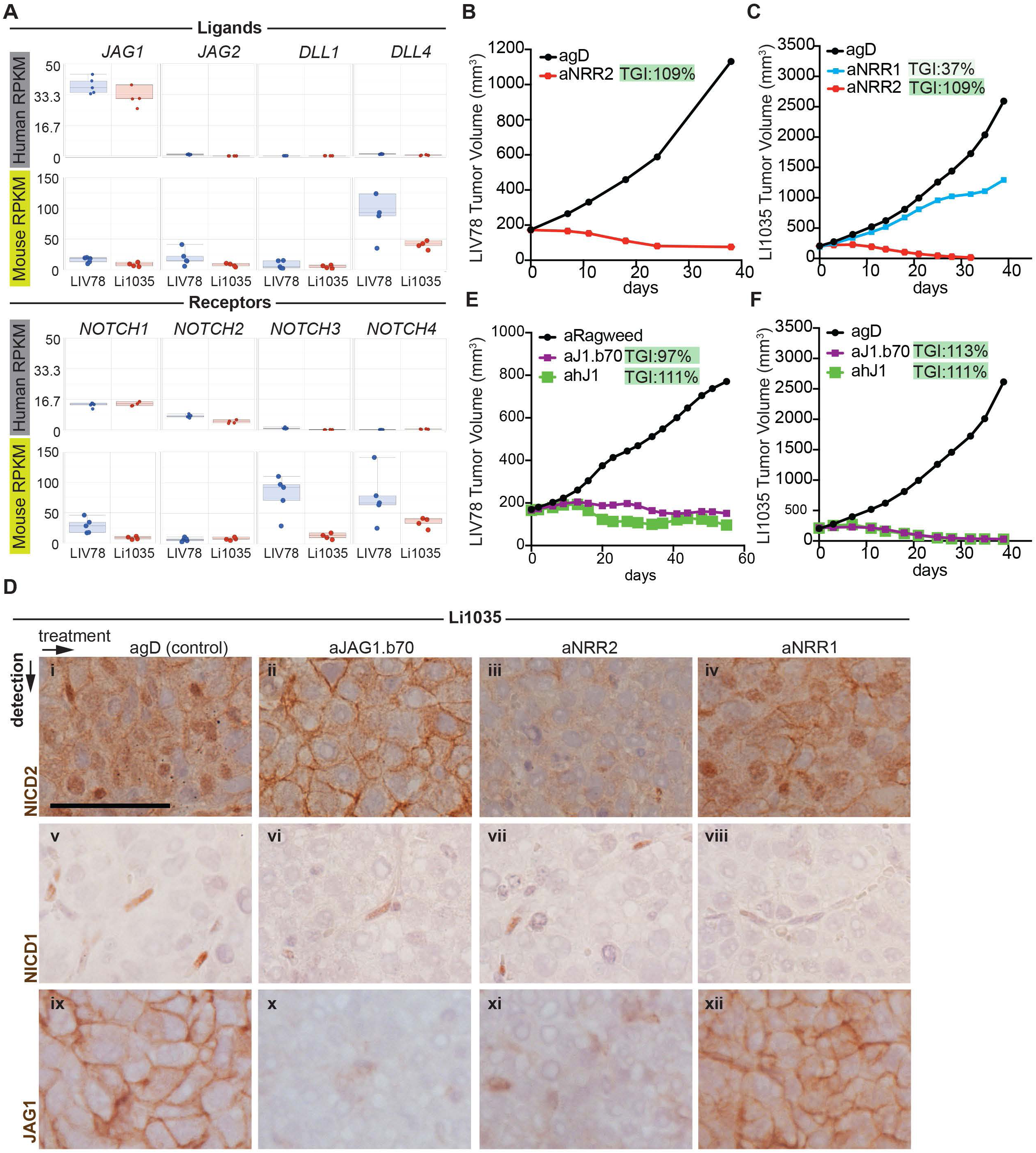
Treatment sensitive tumors depend on an intrinsic JAG1 to NOTCH2 signal **A)** Expression of Notch ligands and receptors in tumor cells (human reads) and stroma (mouse reads) in models LIV78 (blue, n=5) and Li-1035 (red, n=4). Data was generated from tumors 3 days after single dose treatment with control antibody. **B)** Efficacy of aNRR2 treatment in LIV78 model. **C)** Efficacy of aNRR1 or aNRR2 treatment in Li-1035 model. **D)** Immunohistochemical detection of NICD1, NRR2 and JAG1 in model Li1035 treated with control antibodies or antibodies blocking JAG1 or the indicated receptor. Scale bar: 100µm **E)** Efficacy of treatment with aJ1.b70 or ahJAG1 (=aJ1.b6A9, human specific, in LIV78 model. **F)** Efficacy of aJ1.b70 or ahJAG1 treatment in Li-1035 model. Extent of growth response in panels B,C,E, and F is highlighted as in Fig.1B.

To determine the functionally relevant source of JAG1, which could be human JAG1 expressed on the cancer cells or mouse JAG1 expressed on stromal cells, we developed a novel anti-JAG1 blocking antibody. This antibody, aJ1.b6A9, is a mouse anti-human IgG that selectively inhibits human JAG1 but does not inhibit mouse JAG1 (Supplemental Fig. 2D). Treatment with this human-specific antibody induced regressions in both tumor models (Figs. 2E-F, purple line) in the same manner as did aJ1.b70, which cross-reacts with human and mouse JAG1 (Lafkas et al., 2015). Taken together, our results demonstrate that a JAG1-NOTCH2 signaling axis within the human cancer cell population drives the Notch dependency in the super-responder models, with NOTCH2 serving as the functionally relevant receptor.

### Co-expression of cholangiocyte and hepatocyte features distinguish the Notch-dependent HCC tumor models

To better understand the hallmarks of the Notch-dependent tumors, we performed RNA sequencing on all HCC models within the PDX panel and focused on the tumor (human) transcriptomes. K-means analysis of the most variably expressed genes yielded 5 clusters (Fig. 3A), with the sensitive models LIV78 and Li1035 forming a sub-cluster within Cluster 3 (Fig. 3A, B). Comparing the expression signatures of these clusters to cell-type specific signatures generated for normal human fetal and adult liver (Segal et al., 2019) revealed that Cluster 3 is enriched for features of fetal hepatocytes and scores positive for signatures of adult hepatocytes and mixed-lineage hepatobiliary hybrid progenitors (HHyPs) (Segal et al., 2019) (Supp. Fig. 2A).

**Figure 3.**
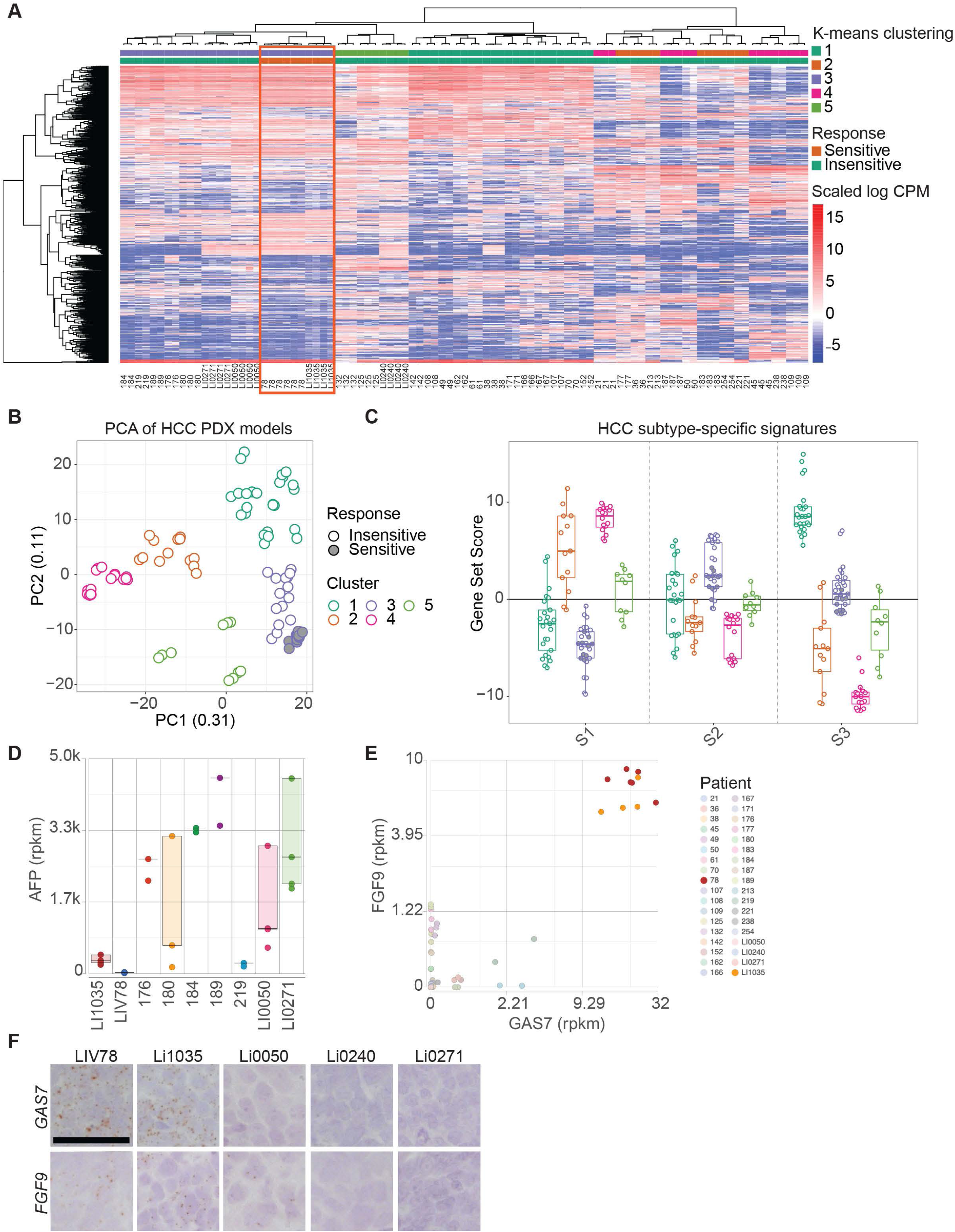
JAG1/NRR2 Inhibition-sensitive HCCs are progenitor-like. **A)** Heatmap of top 1,000 most variable expressed genes in all tested HCC PDX models at baseline (48-72hrs post single dose control antibody treatment). K-means clustering highlights sensitive models as subset of one larger cluster. **B)** PCA of HCC PDX models colored by k-means clustering. **C)** Expression of HCC subtype specific signatures in sensitive and non-sensitive models stratified by k-means cluster (Goossens et al., 2015b). **D)** Expression of AFP in sensitive and insensitive HCC models in k-means cluster 3. **E)** Co-expression of GAS7 and FGF9 separates sensitive and non-sensitive models. **F)** RNAscope detection of GAS7 or FGF9 mRNA expression on tissue sections from treatment responsive or non-responsive tumors. Scale bar: 50µm.

We then focused the gene signature comparison to the S1, S2 and S3 HCC subtypes (Hoshida et al., 2009). S1 and S2 subtypes represent aggressive and poorly/moderately-differentiated tumors, enriched for TP53 mutations and WNT signaling (but not CTNNB1 mutations); S2 tumors typically express AFP, EPCAM, and GPX3, and are marked by IGF2 signaling, a hepatoblastoma-like gene signature, and cancer stem cell features, while S3 tumors are the least aggressive and hepatocyte-like (Goossens et al., 2015a). We found that Cluster 3 was the most S2-like and least S1-like when compared to the other four clusters (Fig. 3C). Although the S2 classification enriched for Notch-dependent models, it was insufficient to fully predict this dependency, as non-responding models also fell within Cluster 3 and the Hoshida S2 type (Fig. 3C). Histological examination of models in Cluster 3 did not reveal any obvious distinguishing features between sensitive and insensitive models also (Supplemental Fig. 2B). However, we noticed that LIV78 and Li1035 express AFP at a low level, atypical for S2 tumors (Fig. 3D). Furthermore, differential gene expression analysis revealed that the Notch-dependent PDX models show a distinguishable upregulated expression of FGF9 and GAS7 (Figs. 3E, F, Supplemental.Table 1), compared to the insensitive models within Cluster 3. In contrast to the PDX dataset, however, FGF9 and GAS7 expression was readily detectable in the majority of patient samples and did not enable subset classifications (Supplemental Fig. 2C). We speculate that FGF9/GAS7 expression in non-tumor cells, absent in our PDX analysis but present in patient samples, may limit the usefulness of these markers as clinical diagnostics. Nevertheless, our results highlight the discovery of an HCC subtype within the aggressive and poorly differentiated S2 class, that exquisitely depends on Notch signaling and resembles a hybrid progenitor cell expressing FGF9 and GAS7.

### Notch inhibition promotes tumor cell differentiation towards the hepatocyte lineage

To elucidate the molecular consequences of Notch inhibition, we performed RNA sequencing of LIV78, Li1035 and three non-responsive models within Cluster 3 (Li0050, Li0240, and Li0271) after treatment with various Notch-blocking and control antibodies. At 72 hours after treatment, JAG1 blockade induced expression changes in hundreds of genes in both the LIV78 and Li1035 tumors; in clear contrast, few or no changes were observed in the three insensitive models (Fig. 4A, Supplemental Fig. 3A, Supplemental Table2). These results likely reflect the presence of an active JAG1-dependent transcriptional program in the regressing models and its absence in the insensitive models. Gene expression changes started as early as 8 hours after treatment, increased in number at 24 and 72 hours, and persisted at least until seven days (Supplemental Fig. 3B, C, Supplemental Table2). NOTCH1 inhibition failed to induce changes in gene expression (Fig. 4A, Supplemental Fig. 3B, Supplemental Table2), consistent with the lack of tumor growth inhibition (Fig. 2C). In contrast, gene expression changes induced by NOTCH2 inhibition were strikingly similar to those caused by JAG1 blockade in both sensitive models (Fig. 4A, Supplemental Fig. 3B, Supplemental Table2), supporting the conclusion that JAG1 and NOTCH2 co-drive the Notch transcriptional program.

**Figure 4.**
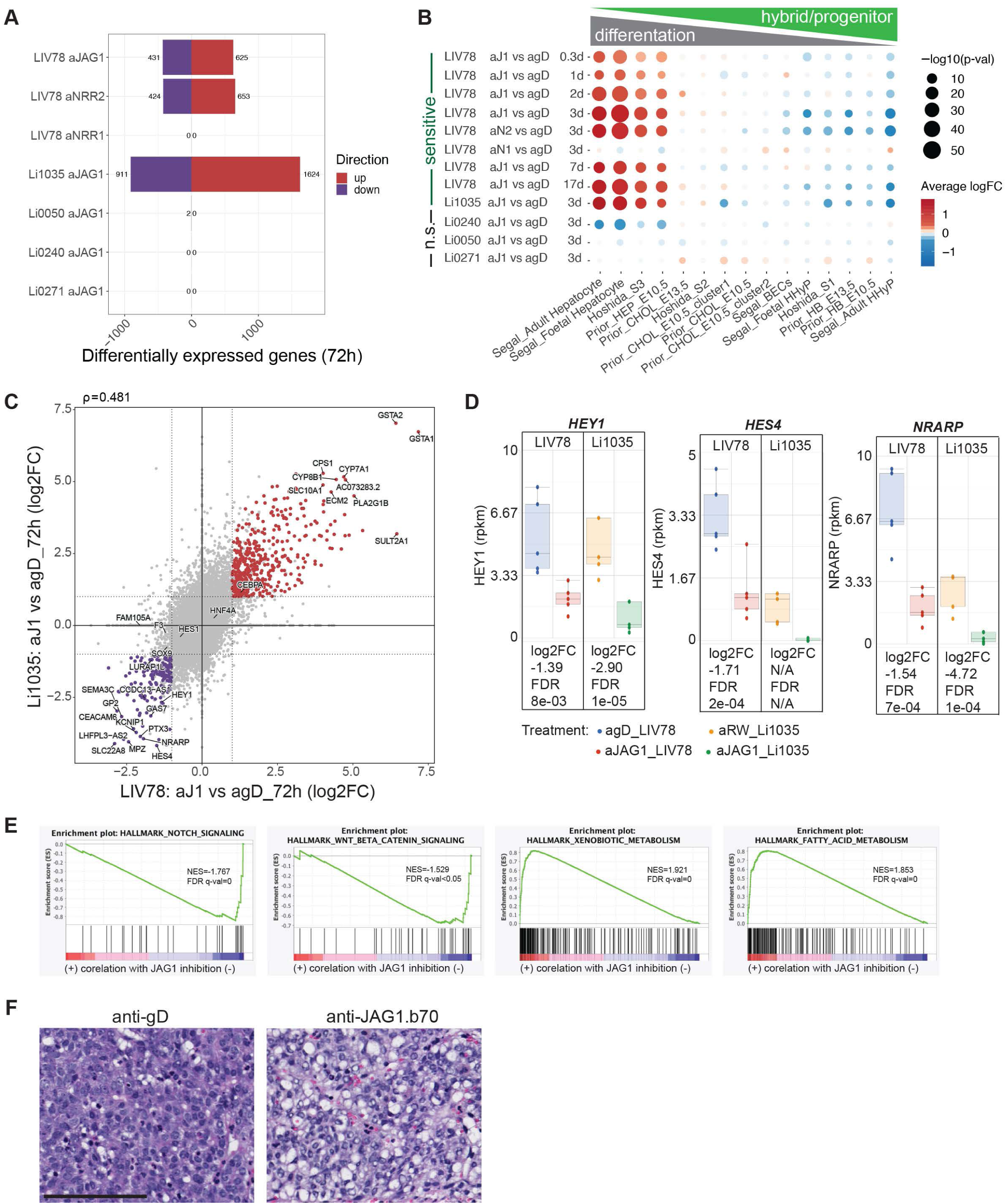
JAG1/NRR2 Inhibition induces progenitor-to-hepatocyte conversion. **A)** Summary of significant gene expression changes following JAG1 inhibition in regressing and non-regressing models. For LIV78 expression changes following NRR1 or NRR2 inhibition are shown. Significantly changed transcripts [log_2_ fold change ≥|1|; FDR<0.05] post single dose treatment are plotted. **B)** Differential expression of cell type or tumor-type specific signatures (Goossens et al., 2015b; Prior et al., 2019;et al., 2019). Changes are obtained from bulk RNA sequencing data for sensitive and insensitive models at 3d post single dose treatment with the indicated blocking antibody. For LIV#078 data is shown for multiple consecutive timepoints in case of treatment with aJ1.b70 vs control antibody. **C)** Four-way comparison of differential gene expression highlights significant overlap in upregulated genes in LIV78 and Li1035. **D)** Expression of NOTCH target genes 72 hrs after JAG1 inhibition in LIV78 and Li1035. N/A: not computed because of low expression values across all groups.**E)** GSEA of differential expression between aJ1.b70 and control antibody treated (72hrs) LIV78 tumors using MSigDB hallmark gene sets. NES–normalized enrichment score, FDR–false discovery ratio. **F)** Hematoxylin and eosin stained LIV78 tumors (17d). Scale bar: 100µm.

To ascertain the consequences of these changes, we mapped the treatment-induced expression signatures onto published signatures of normal liver development and the HCC signatures defined by Hoshida et al. Blockade of JAG1 or NOTCH2 (but not NOTCH1) drove the signatures away from those that describe hybrid/progenitor states towards those of differentiated hepatocytes, with the magnitude and significance of the shifts increasing over the first three days (Fig. 4B). The gene-specific changes underlying this progenitor-to-hepatocyte shift were nearly identical in LIV78 and Li1035 (Fig. 4C) and included decreases in common Notch transcriptional targets (Fig. 4D-E, Supplemental Fig. 3D-F) and increases in hepatocyte metabolic enzymes (Fig. 4C, E). Together, these results indicate that a JAG1-NOTCH2 signaling axis, not involving NOTCH1, drives a shared Notch transcriptional program in the Notch-dependent super responder HCC models.

Gene set enrichment analysis (Liberzon et al., 2015; Subramanian et al., 2005) of the down-regulated transcriptional program not only reinforced the connection to Notch signaling but also highlighted other developmental signaling pathways (‘Wnt beta-catenin signaling’ and ‘TNFa signaling via NFkB’) (Fig. 4E, Supplemental Fig. 3G, Supplemental Table3). JAG1 inhibition also reduced the expression of *SOX4*, *SOX9*, *SPP1*, and HNF1B (Supplemental Fig. 3I-L)---genes that mark cholangiocytes and bi-potent liver progenitors (Coffinier et al., 2002; Lesaffer et al., 2019; Poncy et al., 2015). Similarly down-regulated were gene sets associated with cell proliferation (‘E2F targets’, G2M checkpoint’, ’Myc targets 1 and 2’) (Supplemental Fig. 3G,H, Supplemental Table3). DNA content analysis confirmed that JAG1 inhibition reduced the number of cells in the S and G_2_-M phases (Supplemental Fig. 3M), indicating a treatment-induced cell cycle block in G_0_-G_1_, further supported by reductions in Ki67 staining and BrdU incorporation (Supplemental Fig. 3N,O).

JAG1 inhibition in LIV78 and Li1035 led to increased expression of ‘Hallmark’ gene sets related to ‘xenobiotic metabolism’ or lipid and bile-acid biosynthesis processes (‘Fatty Acid Metabolism’, ‘Peroxisome’, ‘Cholesterol Homeostasis’, ‘Bile acid Metabolism’) (Fig. 4E, Supplemental Fig 3G,H, Supplemental Table3), all specialized functions of mature liver hepatocytes (Stanger, 2015; Trefts et al., 2017). The top upregulated transcripts included genes encoding glutathione transferases (GSTA1-3), essential for enzymatic detoxification of electrophilic compounds such as xenobiotics and oxidative stress products, as well as several members of the cytochrome P450 family, which catalyze endobiotic oxidation and xenobiotic detoxification (Fig. 4C) (Bachmann, 1996; Ramsay & Dilda, 2014). Hepatocytes are also key to regulating lipid homeostasis through de novo synthesis and fatty acid uptake (Trefts et al., 2017). Notch inhibition upregulated genes functioning in fatty acid metabolism, such as the apolipoprotein encoding genes ApoA4, ApoC3, and ApoL1 (Supplemental Fig. 3P-R) and induced an increase in tumor fat content (Fig. 4F). Furthermore, JAG1 blockade induced expression of albumin (ALB), a classic hepatocyte marker (Supplemental Fig. 3S). Taken together, our results support a model in which JAG1-NOTCH2 inhibition targets sensitive HCCs by promoting differentiation of a progenitor/hybrid lineage to a mature hepatocyte phenotype, which is accompanied by cell cycle exit.

### scRNA sequencing analysis reveals heterogeneity of LIV78 tumors and co-expression of JAG1 and NOTCH2

To clarify the cellular composition and the mechanism of efficacy following JAG1/NOTCH2 inhibition, we performed single-cell sequencing of LIV78 tumors isolated from control animals or following inhibition of JAG1 or NOTCH2. Analysis of human cells from all treatment groups yielded six distinct clusters of cells characterized by the expression of markers for: hepatoblasts with high extracellular matrix (ECM) and endoplasmic reticulum (ER) components (cluster 0), oxidative phosphorylation (cluster 1), cell cycle/proliferation (cluster 2), hepatocytes (cluster 3), hypoxia (cluster 4), and ribosomal components (cluster 5) (Fig. 5A). Given the heterogeneity of NOTCH2 activation in the sensitive HCCs (Fig. 2D and Supplemental Fig. 1B) and the separation of ligand- and receptor-expressing populations in other tumor types (Lim et al., 2017), we hypothesized that ligand- and receptor-expressing cells would fall into different clusters. Instead, we found many cells that co-express JAG1 and NOTCH2, with 55% of NOTCH2-positive cells also expressing JAG1 (Supplemental Fig. 4A, B). Ligand and receptor expressing cells co-contributed to multiple clusters, including the largest cluster (0, hepatoblasts), with common Notch targets expressed in a similar distribution (Supplemental Fig. 4C), highlighting cells active for Notch signaling. In contrast, clusters 1 and 5 showed few cells that expressed JAG1 or NOTCH2 (Supplemental Fig. 4A, B). These results suggest that sensitive tumors consist of heterogeneous populations of cells in distinct transcriptional states and that Notch signals emerge from ligand-receptor co-expressing cells.

**Figure 5.**
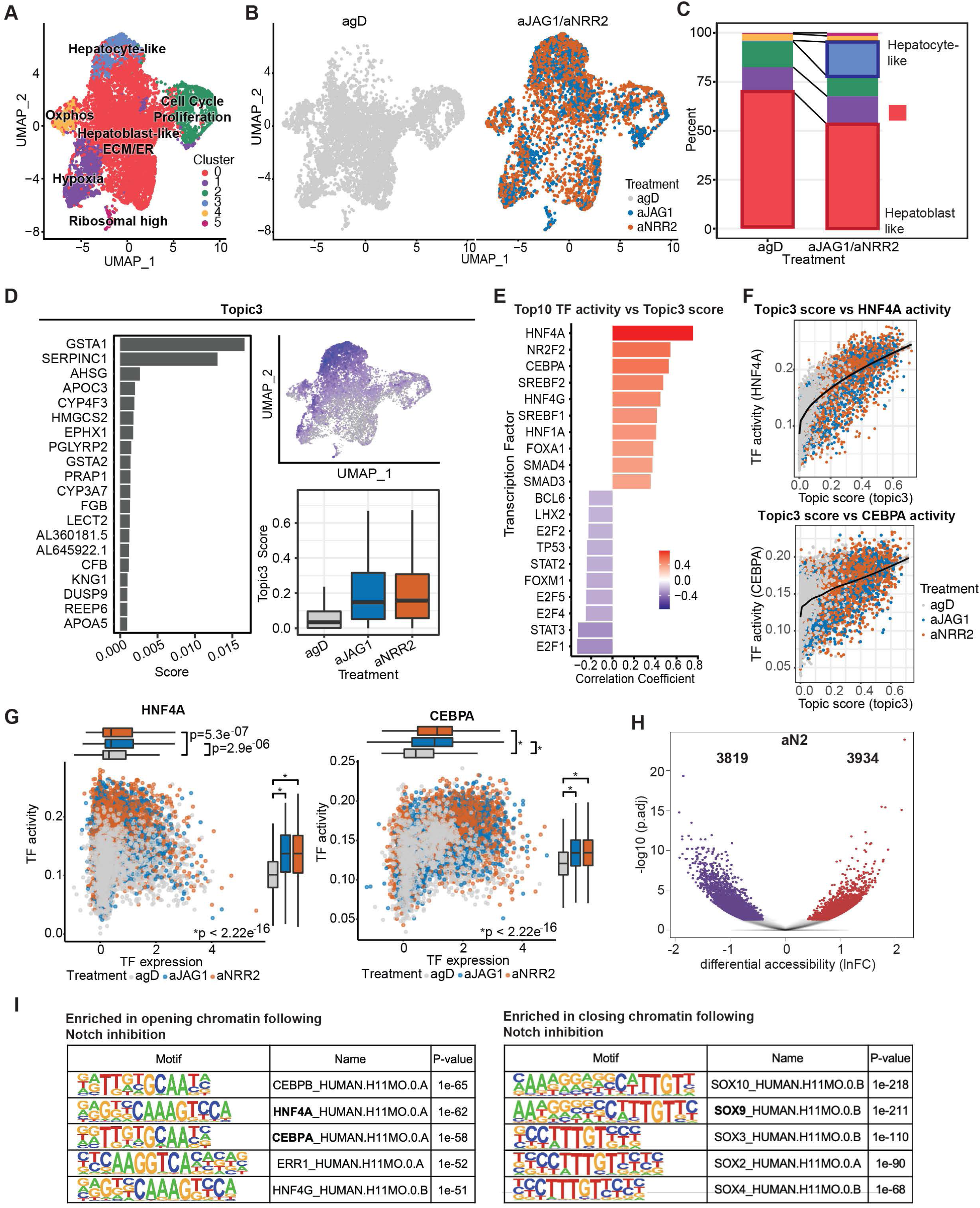
Single cell level analysis identifies HNF4A and CEBPA as Notch controlled transcriptional regulators responsible for progenitor to hepatocyte-like switch underlying treatment efficacy. **A)** UMAP plot of LIV78 tumor cell populations with cluster associations indicated. **B)** UMAP plot of LIV78 tumor cell populations faceted by treatment groups. **C**) Cluster distribution comparison of tumors with Notch signaling inhibition (aJAG1/NOTCH2) and control (agD) tumors. Cl, cluster. **D)** Identification of topic (Topic3) associated with Notch inhibition. Top 20 genes associated with topic3 (left). UMAP color-coded by Topic3 score (top right), topic3 score comparison between treatment groups (bottom right). **E**) Top 10 transcription factor activities that correlate positively or negatively with Topic3 program. **F)** Correlation of Topic3 score and HNF4A activity (top) or CEBPA activity (bottom) with treatment groups indicated. **G)** Correlation of transcription factor expression vs activity for HNF4A (left) and CEBPA (right) with treatment groups indicated. Boxplots provide summary of expression (x-axis) or activity (y-axis) changes with treatment. **H**) Volcano plot of differentially accessible regions in tumors treated with aNRR2 or control antibody (3d). Values with logFC>|1| and adjusted P-value <0.05 are highlighted in red and blue, respectively. **I)** Motif analysis in differentially accessible regions following NRR2 inhibition. The top 3 transcription factors with significantly differential activity values are shown.

### Notch inhibition triggers a cell state change regulated by a CEBPA-HNF4A transcriptional program

Given that Notch signaling commonly regulates binary cell fate decisions (Artavanis-Tsakonas et al., 1999; Koch et al., 2013), including self-renewal versus differentiation of progenitors, we investigated whether treatment affected the distribution of cells between the clusters. Notch blockade dramatically increased the fraction of cells most resembling mature hepatocytes (cluster 3, blue), with 18% of cells from treated tumors contributing to this cluster compared to 0.5% of cells from controls. Concomitantly, JAG1/NOTCH2 blockade reduced the cellular contribution to the hepatoblast-like progenitor population (cluster 0, red) from 70% in controls to 53% after treatment (Fig. 5B, C and Supplemental Fig. 4D-H). This shift from the progenitor to hepatocyte state confirms our bulk RNA-sequencing results (Fig. 4B) and further corroborates our hypothesis that Notch inhibition induces hepatocyte differentiation in the sensitive progenitor-like tumors.

To illuminate the transcriptional programs regulating this progenitor-to-hepatocyte transition, we used Latent Dirichlet Allocation (LDA) to infer continuous transcriptional programs (topics) within cells and correlated these topics with transcription factor (TF) activities for 289 TFs across all cells. Among the nine topics that describe the single-cell data (Supplemental Fig. 4I, J), Topic 3 most strongly associated with cells after treatment (Fig. 5D, Supplemental Fig. 4J). Two of the top three TF activities that best correlated with Topic 3 were HNF4A and CEBPA (Fig. 5E, F), both key regulators of hepatocyte differentiation (Flodby et al., 1996; Watt et al., 2003) that codependently localize on chromatin as cooperatively functioning TF partners (Stefflova et al., 2013). The Topic 3 enrichment for HNF4A activity did not stem from increases in HNF4A protein Supplemental Fig. 4K) or mRNA, the expression of which was only modestly modulated by Notch inhibition and indistinguishable across baseline clusters (Fig. 5G and Supplemental Fig. 4L). In contrast, enrichment for CEBPA activity paralleled increased CEBPA mRNA expression after Notch inhibition (Fig. 5G and Supplemental Fig. 4M), consistent with Notch suppressing CEBPA transcription in other cell types (De Obaldia et al., 2013).

To assess how Notch inhibition impacts chromatin accessibility and how chromatin changes relate to HNF4A and CEBPA activities, we performed Assay-for-Transposase-Accessible-Chromatin (ATAC) sequencing of LIV78 tumors, 72 hours after treatment with NOTCH2 blocking or control antibodies. NOTCH2 blockade profoundly changed the chromatin landscape, increasing or decreasing accessibility at 3934 and 3819 peaks, respectively (Fig. 5H, Supplemental Table 4), with HNF4A and CEBPA binding motifs enriched in chromatin regions with increased accessibility and the SOX family TF motifs in regions exhibiting decreased accessibility (Fig. 5I). HNF4A and CEBPA motifs localized in a characteristic pattern, with HNF4A binding close to the centers of opening regions and CEBPA depleted at these center positions but enriched up- and down-stream (Supplemental Fig. 4N).

We integrated chromatin accessibility data with gene expression measurements, using the BETA software to illuminate how these chromatin changes might relate to differences in gene expression. The increased accessibility of chromatin following NOTCH2 inhibition significantly correlated with increases in gene expression, with the greatest increases in genes related to hepatocyte maturation, which harbor HNF4A and CEPBA motifs in their promoter region. (Supplemental Fig. 4O, Supplemental Table 4). Conversely, decreased accessibility of chromatin correlated with down-regulation of genes associated with progenitor or bile duct cells, which had SOX transcription factor motifs present in the promoter region (Supplemental Fig. 4P, Supplemental Table4). Our TF activity and chromatin analyses thus lead to a model in which Notch inhibition leads to increased expression of CEBPA, thus enabling CEBPA to partner with HNF4A to jointly drive transcriptional programs and the underlying chromatin rearrangements that promote differentiation of progenitor-like tumor cells to a mature hepatocyte fate incompatible with tumor growth and maintenance (Fig. 6).

**Figure 6.**
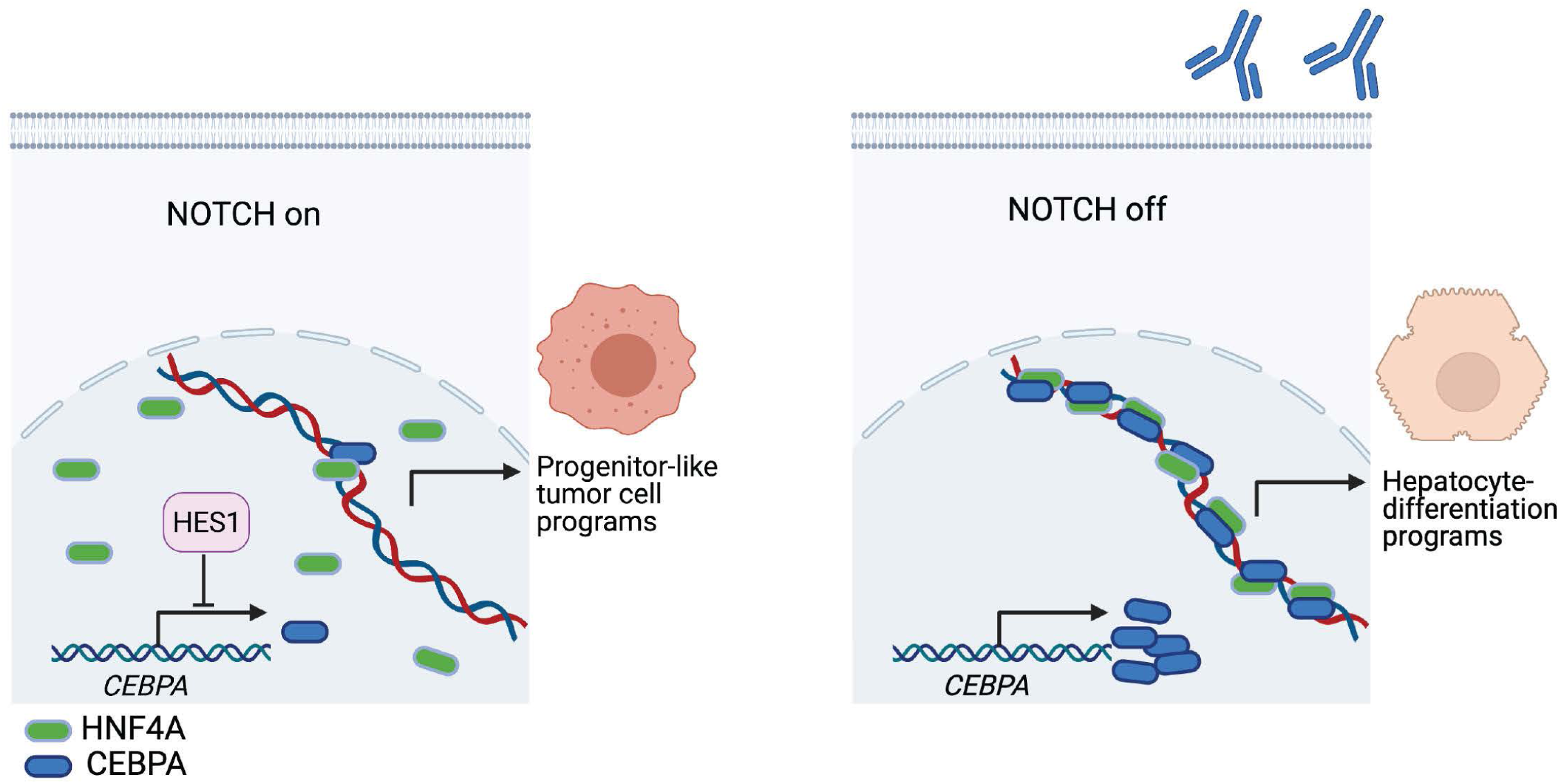
Mechanism of Notch signaling inhibition-induced tumor cell differentiation. In tumor cells with active NOTCH signaling, HES1 is expressed and represses transcription of CEPBA. In absence of CEBPA, HNF4A does not engage with chromatin regions associated with genes involved in hepatocyte differentiation and the program is repressed, locking the tumors cells in a proliferative, progenitor-like state (left panel). Conversely, following inhibition of the relevant JAG1 ligand or NOTCH2 receptor with specific blocking antibodies, HES1 is no longer available to repress *CEBPA* expression. CEBPA becomes available and enables HNF4A’s interaction with chromatin regions of genes functioning during hepatocyte differentiation (right panel). Initiation of hepatocyte differentiation is incompatible with tumor maintenance. Created with <u>BioRender.com</u>

## Discussion

### Discovery of a Notch-dependent, super responder HCC subtype

Our previous work in a liver cancer mouse model (Huntzicker et al., 2015b) prompted us to test the efficacy of selective Notch-targeting therapeutic antibodies in 45 liver tumor PDX models—to our knowledge, the largest *in vivo* liver cancer screen that has been reported. Our studies uncovered an HCC subtype that depends on Notch signaling, and the molecular and cellular mechanisms we describe provide a diagnostic foundation to help identify patient populations predicted to benefit from a Notch inhibitor.

Studies of Notch signaling highlight a complex, multifaceted role in cancer in general and a confusing, contradictory literature in liver cancer in particular. Notch has been described as both oncogenic and tumor suppressive, including in HCC (Dill et al., 2013; Viatour et al., 2011; Villanueva et al., 2012; Zhu et al., 2021). Moreover, studies disagree on the functional relevance of particular ligands and receptors, including NOTCH1 versus NOTCH2 (Dill et al., 2013; Kawaguchi et al., 2016; Villanueva et al., 2012). Some of the conflicting conclusions may reflect choices in model systems, reliance on receptor gain-of-function methods, tumor versus stromal signals, or differences amongst distinct HCC subtypes. The extent of tumor growth inhibition *in vivo* must also be noted. Whereas some dramatic responses to single-agent Notch inhibition have been observed, particularly in tumor models carrying Notch-activating mutations (Agnusdei et al., 2014; Ferrarotto et al., 2017; Stoeck et al., 2014; Y. Wu et al., 2010), regressions or stasis have been rarely achieved, including in our own work testing Notch inhibitors in multiple oncology indications (e.g., Choy et al., 2017). Our current study relies on a solid foundation, built by combining patient-derived xenografts, considered as “high-bar” preclinical models, with therapeutic antibodies highly selective for individual Notch ligands and receptors. While this experimental approach identified numerous models that displayed modest to moderate sensitivity (26-57% TGI), we also identified two exceptional models that were exquisitely sensitive to Notch inhibition. PDX tumors from LIV78 and Li1035 regressed after Notch blockade in a manner that was consistent, reproducible, rapid, and durable, thus displaying a “super responder” phenotype (Marx, 2015; Wheeler et al., 2021) unique to our experience. By using antibodies that selectively block individual ligands and receptors, we identified JAG1 and NOTCH2 as the functionally relevant ligand-receptor pair. While other Notch receptors, particularly NOTCH1 (Villanueva et al., 2012), have been proposed as HCC drivers, we found that NOTCH1 in the super responders was primarily expressed in tumor stromal cells (likely endothelial) and was not required for tumor maintenance. Our results thus highlight the same ligand-receptor pair that determines cholangiocyte versus hepatocyte fate during liver development and reveal the potential of JAG1 and NOTCH2 for targeted therapies in this HCC subclass.

Intriguing questions remain to explain what drives Notch signaling in the super responder subclass. Receptor mutations in the juxtamembrane negative regulatory region (NRR) that enable ligand-independent signaling or in the PEST domain degron that prolong signaling, as well as other genomic alterations that increase Notch signal strength, have been described in multiple cancer types; however, such changes have not been broadly observed in HCC, and we did not find any in our super responders. Similarly, *JAG1* and *NOTCH2* were expressed in the majority of the PDX models and thus were insufficient to mark a Notch-dependent phenotype. However, NOTCH2 activity, defined by nuclear localization of NICD2 and expression of Notch transcriptional targets, was exclusive in the LIV78 and Li1035 tumor cells. JAG1 blockade inhibited this NOTCH2 activity, demonstrating its ligand dependence. In our PDX models, the relevant JAG1 source could be mouse endothelial cells, which express JAG1 (Loomes et al., 1999), or human tumor cells, in which we detected JAG1 mRNA and protein. The newly developed aJ1.b6A9 antibody, which inhibits human but not mouse JAG1, blocked tumor cell NOTCH2 activity and induced regressions. Thus, we conclude that JAG1 induction of NOTCH2 signaling within or between human tumor cells is required for tumor maintenance whereas endothelial (mouse) JAG1 is likely not a major contributor. This tumor cell intrinsic ligand-dependent Notch activation mechanism clearly differs from the mutational activation mechanisms previously described.

Molecular analyses showed that the super responder models display fetal progenitor-like properties and associate with the S2/proliferative HCC subclass. Genetically engineered Notch activation in mouse liver cells positive for AFP, a fetal progenitor marker, was sufficient to drive formation of HCCs (Dill et al., 2013; Villanueva et al., 2012). These Notch-driven mouse tumors display transcriptional profiles that include developmentally-restricted imprinted genes like *Igf2* and *Meg3*, and they classify as undifferentiated and proliferative, thus bearing striking resemblance to LIV78 and Li1035. The proliferative classification, particularly within the Hoshida-S2 subclass, has been previously linked to Notch activation and hepatitis infection (Goossens et al., 2015b and references therein), both of which we note in LIV78 and Li1035. Specifically, signatures derived from human HCCs with Hepatitis C etiology and mouse HCCs driven by Notch clustered together as S2/proliferative (Villanueva et al., 2012). While Notch has also been linked to the S1 subtype (Zhu et al., 2021), this link relied on a GSEA signature predominantly consisting of pathway components with undemonstrated relevance to pathway activity.

In anticipation of possible clinical tests, it is important to consider how to best identify the subset of HCC patients most likely to benefit from JAG1 or NOTCH2 inhibition. Super responder status was rare (∼5%) in our PDX collection. However, PDX models have undergone a powerful biological selection during the xenografting of human tumor samples in mice, and thus the frequency of Notch-dependent super responders in the human HCC population remains to be determined. Nevertheless, our results suggest that expression of fetal progenitor markers and the previously described S2 and proliferative subclassifications provide a valuable initial filter. We also note that some of our PDX models that displayed these traits responded weakly to JAG1 inhibition, revealing limitations to this diagnostic approach. Among our entire PDX panel, including the non-responsive models that transcriptionally co-clustered with the super responders, we also identified *GAS7* and *FGF9* as individual genes that were highly expressed only in the super responders. Furthermore, expression levels of these genes decreased following Notch inhibition, indicating that they are direct or indirect Notch targets. However, *GAS7* and *FGF9* transcript levels were insufficient to clearly stratify HCC patients using large HCC patient datasets. Understanding this discrepancy requires further study, perhaps including spatial methods (e.g., IHC, spatial transcriptomics) to discern GAS7 and FGF9 expression in cells from the tumor, normal liver, stroma, or immune compartment.

### Mechanism of efficacy of NOTCH inhibition in tumors: a replay of cell fate regulation during development?

We found that the efficacy of JAG1 or NOTCH2 inhibition in sensitive hepatocellular carcinomas is based on the transition of a subset of tumor cells from a progenitor-like state to a mature hepatocyte-like phenotype. This trajectory and its regulation closely recapitulate key events during specification of the hepatocyte lineage during embryonic development; in absence of a Notch signal, bi-potent hepatoblasts give rise to hepatocytes, while cholangiocytes are formed only when JAG1-NOTCH2 signaling is active (McCright et al., 2002; Zong et al., 2009 and references therein). Moreover, several lines of evidence highlight the importance of Notch signaling activation in adult hepatocytes for promoting cholangiocyte programs and driving plasticity (Fan et al., 2012; Yimlamai et al., 2014; Zong et al., 2009).

Importantly, our single cell studies enabled in-depth analyses of the NOTCH-regulated molecular interactions that trigger differentiation along the hepatocyte lineage and tumor regressions. These studies identified two closely related tumor cell clusters, one containing progenitor-like cells and one with mature hepatocyte characteristics, which was primarily composed of cells that experienced Notch inhibition. By inferring transcription factor activities, we identified HNF4A and CEBPA as specifically associated with upregulated programs in the treatment-induced hepatocyte-like cluster. Interestingly, both transcription factors are enriched in the liver and essential for hepatocyte formation during embryonic development (Flodby et al., 1996; Parviz et al., 2003; Wang et al., 1995). Moreover, a network comprising both HNF4A, CEBPA, as well as HNF1A, another hepatocyte-lineage expressed transcription factor, and the cholangiocyte-specific factors HNF6 and HNF1B was found to be regulated by NOTCH signaling during lineage commitment of hepatoblasts (Tanimizu & Miyajima, 2004).

*Hnf4a* deficiency leads to aberrant hepatocyte morphology and function, including changes in cell adhesion that disrupt lobule architecture. Deletion of *Cebpa* in mice severely disturbs hepatocyte maturation, with parenchymal cells displaying mixed hepatocyte and biliary epithelial cell characteristics and forming pseudoglandular structures (Flodby et al., 1996). These structures, which notably resemble pseudoglandular hepatocellular carcinoma or tissue undergoing damage-repair, not only co-express markers for both lineages (including HNF4A and SOX9), but also show upregulated expression of *Jag1* and *Notch2* (Akai et al., 2014; Flodby et al., 1996; Yamasaki et al., 2006). Thus, the progenitor-like state we observed in Notch dependent HCC tumor models at baseline is highly similar to the one *Cebpa* loss creates during hepatogenesis.

Our single-cell studies of the LIV78 model showed that *HNF4A* transcript was expressed across all clusters, with transcript levels showing little to no treatment-dependent variation. In contrast, activity and expression of *CEBPA* strongly increased after Notch inhibition, with particularly high levels in cells contributing to a hepatocyte-like cluster that emerged following Notch-inhibition. Interestingly, CEBPA was shown to be negatively regulated by HES1(a bona fide Notch target gene) (De Obaldia et al., 2013) in the context of myeloid fate regulation. In chronic myelomonocytic leukemia, which results from an inactivating mutation in the y-secretase complex component nicastrin, Notch signaling has a tumor suppressive function, that is mediated through direct repression of CEBPA and PU.1 by HES1 (Klinakis et al., 2011). In the liver, HNF4A and CEBPA were shown to extensively co-localize on chromatin and loss of HNF4A reduces the ability of CEBPA to bind DNA and vice versa (Stefflova et al., 2013), rendering HNF4A necessary, but not sufficient for the development of mature hepatocyte features. The hypothesis that Notch inhibition through de-repressing *CEBPA* transcription allows chromatin engagement of both CEBPA and HNF4A and thus initiation of hepatocyte differentiation programs, is strongly supported by our ATAC sequencing data; chromatin regions that open following NOTCH2 inhibition are strongly enriched for CEBPA and HNF4A motifs which were found in close proximity of each other. Closing chromatin regions on the other hand are associated with motif enrichment for SOX family transcription factors, and HNF1B, re-emphasizing the role of active Notch signaling in maintaining cholangiocyte specific programs active and the progenitor-like tumor cells in a multipotent state.

Our study provides proof-of-concept for a clinical paradigm using therapeutic antibodies targeting JAG1 or NOTCH2 to promote a hepatocyte-like state incompatible with tumor maintenance, and identifies the key regulatory network underlying Notch inhibition-induced tumor cell differentiation. This, together with the finding that therapeutic antibodies against NOTCH2 are well tolerated in patients (Smith et al., 2019), holds great promise for the treatment of Notch-active HCCs and possibly other progenitor-like cancers.

## Material and Methods

### Treatment and tumor growth assessment of liver cancer PDX models

Animals were maintained in accordance with the Guide for the Care and Use of Laboratory Animals (National Research Council 2011). Genentech is an AAALAC-accredited facility and all animal activities in this research study were conducted under protocols approved by the Genentech Institutional Animal Care and Use Committee (IACUC). Mice were housed in individually ventilated cages within animal rooms maintained on a 14:10-hour, light: dark cycle. Animal rooms were temperature and humidity-controlled, between 68 to 79°F (20.0 to 26.1°C) and 30 to 70% respectively, with 10 to 15 room air exchanges per hour.

To generate tumor-bearing animals for the mouse clinical trial and all subsequent studies, tumor fragments from 42 established PDX liver cancer models (comprised of 35 HCC models plus CCs and other liver tumor subtypes) were passaged from tumor-bearing animals and implanted sub-cutaneously (s.c.) into the right flank of either Balb/c nude (NU/J *Foxn1^nu^* (JAX:002019), Jackson Laboratories) or NCr nude (CrTac:NCr-*Foxn1^nu^*;Taconic Biosciences) immunocompromised mice to generate patient-derived xenografts (PDXs). Tumors were allowed to establish until they had engrafted and demonstrated consistent growth (typically between 80-280 mm^3^). Animals were then grouped into treatment cohorts (between n=5-10 across studies) to normalize across groups based upon tumor volume prior to initiation of treatment. During the mouse clinical trial, mice were treated with either IgG1 isotype anti-Ragweed control antibodies or anti-Jag1 (aJ1.b70) antibodies intraperitoneally (i.p.) at 10 mg/kg, weekly (QW) for 3 weeks.

For subsequent studies, either IgG1 isotype anti-gD, anti-Jag1 (aJ1.b70), anti-huma-Jag1 (ahJ1), anti-Notch1 (anti-NRR1), or anti-Notch2 (anti-NRR2) antibodies were used at the indicated doses (between 1.25 to 30 mg/kg), QW, i.p. either for efficacy assessment or generation of samples for analysis of pharmacodynamics. Animals were monitored routinely to assess health and body condition with tumor volumes and body weights being measured twice weekly. Animals demonstrating excessive body weight loss (>15%) or tumors in excess of 1500 mm3 or bearing ulcerations were routinely euthanized. In general all treatments were well-tolerated with the exception of anti-Notch1 (aNRR1), which has known and expected side-effects resulting in body weight loss.

Analyses and comparisons of tumor growth were performed using a package of customized functions in R (Version 3.4.2; R Foundation for Statistical Computing; Vienna, Austria), which integrates software from open source packages (e.g., lme4, mgcv, gamm4, multcomp, settings, and plyr) and several packages from tidyverse (e.g., magrittr, dplyr, tidyr, and ggplot2) (Forrest et al., 2020). Briefly, as tumors generally exhibit exponential growth, tumor volumes were subjected to natural log transformation before analysis. All raw tumor volume measurements less than 8 mm^3^ were judged to reflect complete tumor absence and were converted to 8 mm^3^ prior to natural log transformation. Additionally, all raw tumor volume measurements less than 16 mm^3^ were considered miniscule tumors too small to be measured accurately and were converted to 16 mm^3^ prior to natural log transformation. The same generalized additive mixed model (GAMM) was then applied to fit the temporal profile of the log-transformed tumor volumes in all study groups with regression splines and automatically generated spline bases. This approach addresses both repeated measurements from the same study subjects and moderate dropouts before the end of the study.

Estimates of group-level efficacy were obtained using the calculation of percent tumor growth inhibition (TGI). This value represents the percent difference in tumor growth between the treatment and reference groups according to each group fit baseline-corrected area under the curve (AUC). This AUC is obtained after back-transforming tumor volumes to the original scale and correcting for starting tumor burden. Positive values indicate antitumor effects, with 100% denoting stasis and values >100% denoting regression (negative values indicate a pro-tumor effect). The 95% confidence intervals are based on the fitted model and variability measures of the data obtained using parametric bootstrapping.

### Generation and characterization of human-specific anti-Jagged1 antibody

For development of anti-huJagged1 hybridoma, five Balb/c mice (Charles River Laboratories International, Inc., Wilmington, MA, USA) were hyperimmunized, in each hind footpad at 3-4 day intervals, with recombinant soluble human Jagged1 ECD in RIBI adjuvant (Ribi Immunochem Research Inc. Hamilton, MT, USA). After 5 injections, serum titers were evaluated by standard enzyme-linked immunosorbant assay (ELISA) to identify mice with positive serum titers to huJagged1 ECD. B cells from popliteal lymph nodes from five mice were pooled and were fused with mouse myeloma cells (X63.Ag8.653; American Type Culture Collection, Manassas, VA, USA) by electrofusion (ECM 2001; Harvard Apparatus, Inc., Holliston, MA, USA). After 10-14 days, hybridoma supernatants were harvested and screened for antibody production with a huJagged1 binding ELISA. All specific clones were then re-screened for binding by FACS (293/3T3-huJagged1 cells) and for blocking Notch1-Jagged1 signaling by co-culture assay. Hybridoma clone, 6A9.7.3, showed high immunobinding and signal blocking after the second round of single cell per well subcloning (FACSAria cell sorter; BD Biosciences, San Jose, CA, USA) and was scaled up for purification in INTEGRA CELLine 1000 bioreactor (INTEGRA Biosciences AG, Zizers, Switzerland). Supernatant was purified by protein A affinity chromatography, sterile-filtered, and stored at 4°C in PBS. Isotype of the mAb was determined using the Isostrip Mouse mAb Isotyping Kit (Roche Applied Biosciences, Indianapolis, IN, USA). The isotype was determined to be IgG1, kappa. Clone 6A9.7.3 was sequenced and chimerized into huIgG1, kappa backbone.

Notch reporter assays were performed as previously described (Lafkas et al., 2015) using U87glioblastoma cells co-transfected with a Notch-response TP-1 (12XCSL) firefly luciferase reporter and a constitutively expressed *Renilla* luciferase reporter (pRL-CMV, Promega E2261). Antibodies were added with the ligand-expressing cells (NIH-3T3 cells stably transfected with human *JAG1* or OP9 cells stably transfected with mouse *Jag1*).

### Generation and analysis of bulk RNAseq data

Tumor tissue was harvested, snap frozen in liquid nitrogen and stored at -80C. Total RNA was extracted from 15-25mg intact or powderized tumor tissue using TRIzol^TM^ Reagent (Thermo Fisher Scientific Cat# 15596026) according to the manufacturer’s instructions. Samples were homogenized using a TissueLyser II (Qiagen) at 25k rpm for 3min. The optional 5min incubation to allow for complete dissociation of the nucleoprotein complex was performed. Following isopropanol precipitation, samples were loaded onto RNEasy Mini prep columns (Qiagen) and cleanup steps, including on-column DNAse digestion, were performed according to the RNEasy Kit’s instructions. Samples were eluted in 30ul RNAse-free water.

2-6 replicate samples were collected for each treatment condition. The concentration of RNA was determined using NanoDrop 8000 (Thermo Scientific) and the integrity of RNA was determined by Fragment Analyzer (Advanced Analytical Technologies). Approximately 500 ng of total RNA was used as an input for library preparation using TruSeq RNA Sample Preparation Kit v2 (Illumina). Size of the libraries was confirmed using High Sensitivity D1K screen tape (Agilent Technologies) and their concentration was determined by qPCR-based method using Library quantification kit (KAPA). The libraries were multiplexed and sequenced on Illumina HiSeq 2500 (Illumina). We obtained on average 30 million single-end RNA-seq reads (50bp) per sample.

Sequencing reads were processed using a XenoFilter pipeline to obtain human- and mouse-specific gene expression quantifications. Reads were first aligned to ribosomal RNA sequences to remove ribosomal reads. The remaining reads were aligned to the human reference genome (NCBI Build 38) using GSNAP (T. D. Wu & Nacu, 2010) version ‘2013-10-10’, allowing a maximum of two mismatches per 50 base pair sequence (parameters: ‘-M 2 -n 10 -B 2 -i 1 -N 1 -w 200000 -E 1 --pairmax-rna=200000 --clip-overlap’). The same procedure was used to map all reads to the mouse reference genome (NCBI Build 38). Reads that were multi-mappers in either of the two organisms were removed. Subsequently, concordant and unique reads with <= 3 mismatches to the human genome and >3 mismatches to the mouse genome were called human-specific. The reverse mismatch criteria were used to determine mouse-specific reads. Lastly, gene expression levels were quantified by calculating the number of reads mapped to the exons of each RefSeq gene using the HTSeqGenie R package. Transcript annotation was based on the RefSeq database NCBI Annotation Releases 104 and 106 for mouse and human, respectively. Read counts were scaled by library size, quantile normalized and precision weights calculated using the “voom” R package (Law et al., 2014). Subsequently, differential expression analysis on the normalized count data was performed using the “limma” R package (Ritchie et al., 2015) by contrasting anti-JAG1 or anti-Notch antibody treated samples with control samples. Gene expression was considered significantly different across groups if we observed an |log2-fold change| ≥ 1 (estimated from the model coefficients) associated with an FDR adjusted P-value ≤ 0.05. In addition, gene expression was obtained in the form of normalized Reads Per Kilobase gene model per Million total reads (nRPKM) as described previously (Srinivasan et al., 2016). Graphs were generated in the R environment or using Partek Software.

TCGA gene expression was obtained as raw counts that were estimated with the PanCanAtlas pipeline (https://docs.gdc.cancer.gov/Data/Bioinformatics_Pipelines/ Expression_mRNA_Pipeline) using the human genome reference (NCBI Build 38) and Gencode (V22) gene models. Subsequently, read counts were scaled by library size, quantile normalized and precision weights calculated in the same fashion as for our in-house data (above). PCA was performed on the normalized data and UMAPs were produced using the first 1000 principal components.

### Histology, immunohistochemistry and in situ RNA hybridization

Tumors were isolated, fixed in 10% neutral buffered formalin overnight, washed in two changes of ddH_2_O and dehydrated through a series of increasing concentrations of ethanol and xylenes for paraffin embedding. Tissue sections were prepared at 5μm thickness and slides were incubated at 60C for 10 minutes, de-paraffinized using Xylenes and rehydrated through an ethanol gradient prior to any staining procedure.

For evaluation of morphology, sections were stained with Hematoxylin and Eosin (Fisher Scientific, # 23245637 and #6766010) using standard methods.

For immunohistochemical (IHC) and immunofluorescence (IF) staining, sections were boiled for 15 min in Target Retrieval solution (DAKO-S2369) using a pressure cooker, treated with 3% hydrogen peroxide (VWR-BDH7690-1) for 10 min, washed in PBS, and subjected to a 1 h blocking step in 5% bovine serum albumin (Sigma-A9647) in PBT (PBS with 0.1% Triton-X, Sigma-T9284). For IHC, incubation with primary antibodies from Cell Signaling Technologies against N1-ICD (Val1744, #4147), NOTCH2 (D76A6, #5732), or JAGGED1 (D47Y1R, #70109) diluted 1:2000 in blocking solution was performed overnight in a humidified chamber. Three 10 min washes in PBS were followed by incubation with Poly-HRP anti Rabbit IgG secondary antibody (PowerVision, PV6119) for one hour, two washes in PBS and one wash in PBT, each for 10 min and application of DAB substrate (25μl/2ml buffer, DAB+ Substrate Chromogen system, DAKO-K3468). Staining was performed in the dark for 2 minutes after which the slides were immediately immersed into tap water. To visualize the nuclei, the sections were counterstained for 40 seconds using Gill’s hematoxylin III diluted 1:5 in water. A brief bluing step in Scotts tap water substitute (Eng Scientific, #8442) was included prior to dehydration through ethanols and xylenes and slides were mounted in Permount (Fisher Scientific, #SP15).

For IF staining, sections were incubated in a 1:100 dilution of anti-BrdU antibody (Biorad, MCA2060), 1:100 dilution of anti-HNF4A (Perseus Proteomics, PP-H1415-00), or 1:1000 dilution of anti-Ki67 primary antibody (Thermo Scientific, RM-9106-50). Alexa Fluor 555 goat anti-rat or goat anti-rabbit, or Alexa Fluor 488 goat anti mouse secondary antibodies were used at a 1:250 dilution (Thermo Scientific A-21428, A-21434, and A-11001). Following nuclear staining with DAPI (Invitrogen, D1306), sections were mounted in Pro Long Gold antifade reagent (Invitrogen, P36934).

For in situ RNA hybridization, mRNAs were visualized by RNAscope® technology (Wang et al., 2012). Manual assays were performed using the RNAscope® 2.5 HD Reagent Kit-BROWN (Advanced Cell Diagnostics, #322300) according to the manufacturer’s instructions. Probes against *FGF9* (Cat. No.439431) or *GAS7* (custom order) as well as positive and negative control probes (310451, 310043) were designed by Advanced Cell diagnostics.

All tissue stainings were performed for at least 3 animals per treatment group.

Images were obtained on a Zeiss Axioscope 40 or Nanozoomer slide scanner.

### Quantification of Immunofluorescence staining data

For quantification of BrdU+ or Ki67+ cells, 3 randomly chosen 20x frames were imaged per tumor. Image J software was used to count DAPI stained or marker-stained nuclei. First, images were converted to 8-bit format, a threshold was set and the “watershed” option was used to separate clustered nuclei. The “Analyze Particle” command was used with ‘‘size = 0.005-∝ inch 2’’ and ‘‘circularity = 0.00-∝ ”. Cell counts reported by ImageJ were expressed as % of marker-positive nuclei/DAPI positive nuclei. Graphs were generated using PRISM software. Each data point graphed is the average value from one mouse; data are presented as mean ± SEM.

### Analysis of cell cycle phase by flow cytometry

PDX tumors were harvested, dissociated into single cells and subjected to depletion of CD45 and Ter19-postive populations as described above. Following two washes in ice-cold PBS, each sample was resuspended in 1ml ice-cold PBS and 3ml of ice-cold 100% ethanol were added to the tube, drop-wise while vortexing on slow speed to avoid clumping while fixing the cells. For proprio iodine staining, samples were pelleted, the ethanol removed, and following 2 washes with ice-cold PBS the cells were resuspended in 0.5ml PI/RNAse staining solution (BD550825) per 1x10^6 cells. After passing the cells through a 35um nylon mesh-strainer cap into FACS tubes (Corning Catalog #352235) they were incubated for 15 minutes at RT prior to analyzing on a BD Symphony Analyzer.

### Generation, processing and analysis of single-cell RNAseq data

NCR nude mice bearing similarly sized LIV78 tumors (volumes between 300-900mm^3^) were injected intravenously with 30mg/kg bodyweight of either anti-Notch2 or anti-Jag1 or anti-gD control antibody 72hrs prior to tumor harvest. 2 samples (one with Notch inhibition and one with control antibody treatment) were processed in the same day.

The tumor cell isolation procedure was adapted from a liver cell isolation technique by Mederacke et al (Mederacke et al., 2015). Tumor-bearing mice were sacrificed and the tumors isolated and placed in a petri-dish containing 5mls of a total of 50ml of enzyme buffer solution (8000mg/L NaCl, 400mg/L KCl, 88.17mg/L NaH_2_PO_4_*H_2_O, 2380mg/L

HEPES, 350mg/L NaHCO_3_, 560mg/L CaCl_2_ *2H_2_O) with 25mg of Pronase (Sigma-Aldrich #P5147), 4.4 U of Collagenase D (Roche #11088882001), 32μM Actinomycin D (Chemodex #A0043) and 1% DNAse I (Roche #10104159001). Tumors were minced to small pieces with a razor blade, added to the remaining 45mls of Pronase/Collagenase/ DNAse/Actinomycin D solution in a 50 ml conical tube and digested for ∼1hr rotating in a hybridization oven at 42C. The digested tumors were filtered through a 70μm cell strainer into a new 50ml tube and centrifuged at 450xg for 10min at 4C. Following aspiration of all but 10ml of the supernatant, 120ul of DNAse I stock solution (100mg DNAse I in 50 ml Gey’s Balanced Salt Solution B (GBSS/B; Sigma-Aldrich #G9779)) were added and the cells were resuspended. The tube was filled up to 50ml with GBSS/B to wash the cells prior to centrifugation at 450xg for 10 min at 4C. Tumor cells were resuspended in 3% heat inactivated fetal bovine serum (Sigma-Aldrich #19G057-AH1) in PBS and subjected to removal of host immune and red blood cells using the EasySep^TM^ Mouse Mesenchymal Stem/Progenitor Cell Enrichment kit (Stem Cell Technologies #19771) according to the manufacturer’s instructions.

Following the depletion of CD45+ and Ter119+ cells, the samples were processed for single-cell RNA-seq (scRNAseq) as described previously (Long et al., 2019) using the Chromium Single Cell 3’ Library and Gel bead kit v2, following the manufacturer’s manual (CG00052 Chromium Single Cell 3 Reagent Kits v2 User Guide RevA; 10x Genomics). Cell density and viability of the single-cell suspensions were determined by Vi-CELL XR cell counter (Beckman Coulter). All of the processed samples had more than 95% viable cells. Cell density was used to impute the volume of single-cell suspension needed in the RT master mix. cDNAs and libraries were prepared following the manufacturer’s manual (10X Genomics). Libraries were profiled by Bioanalyzer High Sensitivity DNA kit (Agilent Technologies) and quantified using Kapa Library Quantification Kit (Kapa Biosystems, Wilmington, MA). Each library was sequenced in one lane of HiSeq 2500 (Illumina) following the manufacturer’s sequencing specification (10X Genomics).

Sequencing data were processed as described previously (Long et al., 2019). In short, reads were tallied and demultiplexed based on their association with cell-specific barcodes. Only barcodes with >10K associated reads were considered for further processing. For each cell, reads were mapped to the human and mouse reference genomes using the same GSNAP settings as for bulk RNA, respectively. The number of transcripts per gene was quantified based on unique molecular identifiers (UMI) using reads that overlapped exonic regions in sense direction. Subsequently, cells were classified as human (PDX) or mouse (stroma) if their mapping rate was higher than 42% to the respective genome. Only cells with a mitochondrial UMI fraction <0.25 were considered for analysis. Per-gene UMI counts were normalized by the total number of transcripts per cell and scaled by the median number of transcripts across all cells. Dataset alignment (between multiple processing dates), cell clustering, visualization, calculation of gene set scores and differential expression analysis were performed according to best practices using Seurat (v3.0.0.9000) (Stuart et al., 2019). Differential expression between treatment conditions was based on the likelihood-ratio test for single cell gene expression (McDavid et al., 2013).

To characterize cells based on the activity of transcriptional programs, we performed topic modeling using functions from the CountClust R package (Dey et al., 2017). We evaluated a range between 2 to 15 topics for their fit using the *FitGoM* function. A total of 9 topics were selected to represent the data after inspecting the AIC and BIC statistics. Top genes associated with each topic were identified by applying the *ExtractTopFeatures* function, which uses the relative gene expression profile of the GoM clusters and applies a KL-divergence based method to obtain a list of top features that drive each of the clusters.

To identify transcription factors (TFs) that drive specific transcriptional programs (characterized by individual topics) we quantified the transcription factor activities in individual cells and correlated them with the previously obtained topic weights. We used a total of 289 high-confidence regulons (comprised of TF downstream target genes) from the non-academic version of the Dorothea database (Garcia-Alonso et al., 2019; Holland, Szalai, et al., 2020; Holland, Tanevski, et al., 2020) and quantified their activity using AUcell (Aibar et al., 2017).

To perform differential abundance analysis of cell groups between treatment conditions, we used the MiloR package (Dann et al., 2022). In brief, milo uses a k-nearest neighbor (KNN) graph to define a set of representative neighborhoods, followed by counting of cells in each neighborhood by experimental condition. These counts are used for differential abundance testing using a negative binomial GLM framework followed by a weighted FDR procedure that takes the spatial overlap of neighborhoods into account.

Graphs were generated in the R environment or using Partek Software.

### Gene set enrichment

We tested for the enrichment of particular gene categories to identify relevant biological processes associated with (1) differential expression or (2) transcriptional programs (topics), by using functions from the "clusterProfiler" R package (T. Wu et al., 2021) and the MSigDB gene set collection (Liberzon et al., 2011). In the case of differential expression, treated samples were contrasted with control samples, genes were ranked based on the π statistic (π=logFC * -log10(p-val)) (Xiao et al., 2014) and subsequently gene set enrichment was performed using the *clusterProfiler::GSEA* function or the GSEA4.1.0 software. In the case of transcriptional programs, we extracted the top 50 genes associated with specific topics and performed a hypergeometric test using the *clusterProfiler::enricher* function.

### Generation and analysis of bulk ATAC-seq data

Tumors were isolated, snap frozen in liquid nitrogen and stored at -80C. Tumor tissue was pulverized using a Covaris CP02 CryoPrep Pulverizer. Flash frozen pulverized tumor samples were sent to Active Motif to perform the ATAC-seq assay.

The tissues were manually disassociated, isolated nuclei were quantified using a hemocytometer, and 100,000 nuclei were tagmented as previously described (Buenrostro et al. 2013), with some modifications based on (Corces et al. 2017) using the enzyme and buffer provided in the ATAC-Seq Kit (Active Motif). Tagmented DNA was then purified using the MinElute PCR purification kit (Qiagen), amplified with 10 cycles of PCR, and purified using Agencourt AMPure SPRI beads (Beckman Coulter). Resulting material was quantified using the KAPA Library Quantification Kit for Illumina platforms (KAPA Biosystems), and sequenced with PE42 sequencing on the NextSeq 500 sequencer (Illumina).

The ATAC-seq data were aligned to human reference genome (NCBI Build 38) using the ENCODE ATAC-seq pipeline (https://github.com/ENCODE-DCC/atac-seq-pipeline) with minimum p-value cutoff for peak calling set to 1e-6. The resulting peak lists were sorted and overlapping peaks were merged using bedtools (v2.26.0). The differential accessibility analysis was performed using the bioconductor package DiffBind (v3.2) in R (v4.1.0) (Ross-Innes et al., 2012; Stark & Brown, 2011). For differential accessibility analysis, peaks called in only one sample were filtered out. The read counts in the peak matrix were normalized using the dba.normalize() function of DiffBind with the parameter library = DBA_LIBSIZE_PEAKREADS. The statistical significance of differential accessible regions was estimated using the dba.analyze() function of DiffBind in its default setting. The motif enrichment analysis was performed using the AME (v5.4.1) tool of the MEME suite using the non-differential ATAC-seq peaks as background and HOCOMOCOv11_core_HUMAN motif database (Kulakovskiy et al., 2018; McLeay & Bailey, 2010). The motif localization analysis was performed using the CENTRIMO (v5.3.0) tool from the MEME suite (Bailey & MacHanick, 2012).

## Acknowledgements

We wish to thank Nayab Adibi, Simona Hankeova, and Gloria Hernandez for helpful discussions. We are grateful to Heinrich Jasper, Louis Vermeulen, and Xin Ye for critical reading of the manuscript. We would like to thank the Genentech Necropsy, Histology and Pathology and Legal teams for their contributions and advise.

## Author contributions

K.S., R.P. and C.W.S. designed the study; C.W.S. supervised and directed the project. K.S. performed the experiments and data analysis; R.P., T.T.T. N. and M.A. processed, analyzed and interpreted sequencing data; C.C., A.Sh. and J.C. contributed to experiments and data analysis; J.G. performed in house PDX model studies; C.D.L.C, U.S., and M.M facilitated PDX model studies in house and at CROs and performed data analysis; Y.A.Y., J.C. and Z.M. performed RNA sequencing experiments; A.Sc. provided histopathological analysis and discussion; R.V. and Y.W. generated and validated antibodies used in this study; L.M. provided input for experiments, discussion and edits for the manuscript. C.M. performed experiments and data analysis and provided edits and critical input throughout this study; K.S., R.P., and C.W.S. cowrote the paper. All authors contributed to the manuscript and approve its content.

## Conflict of interest

The authors are or were employed by Genentech Inc., a member of the Roche family of companies, and have received or currently receive stock-based compensation from Roche. The work was funded by Genentech, Inc.

Some of the co-authors are inventors on the following relevant (to the current work) patents and patent applications:

US-20240034805-A1, US-11702479-B2, US-20210206871-A1, US-10858440-B2, US-20200377610-A1, US-10689455-B2,US-20190135936-A1, US-10266602-B2, US-10208114-B2, US-20180371098-A1, US-20180355054-A1, US-10113002-B2, US-10011661-B2, US-10005844-B2, US-9982058-B2, US-20180112005-A1, US-9914774-B2, US-20180057589-A1, US-20170327574-A1, US-20170233486-A1, US-20170210815-A1, US-20170204175-A9, US-20170137531-A1, US-9550829-B2, US-9533042-B2, US-9518121-B2, US-20150252117-A1, US-20150232568-A1, US-20150104461-A1, US-20140314782-A1, US-20140314749-A1, US-20140296488-A1, US-20140271624-A1, US-20140037643-A1.

## Supplemental Figures

**Supplemental Figure 1 (goes with Main Figure2).**
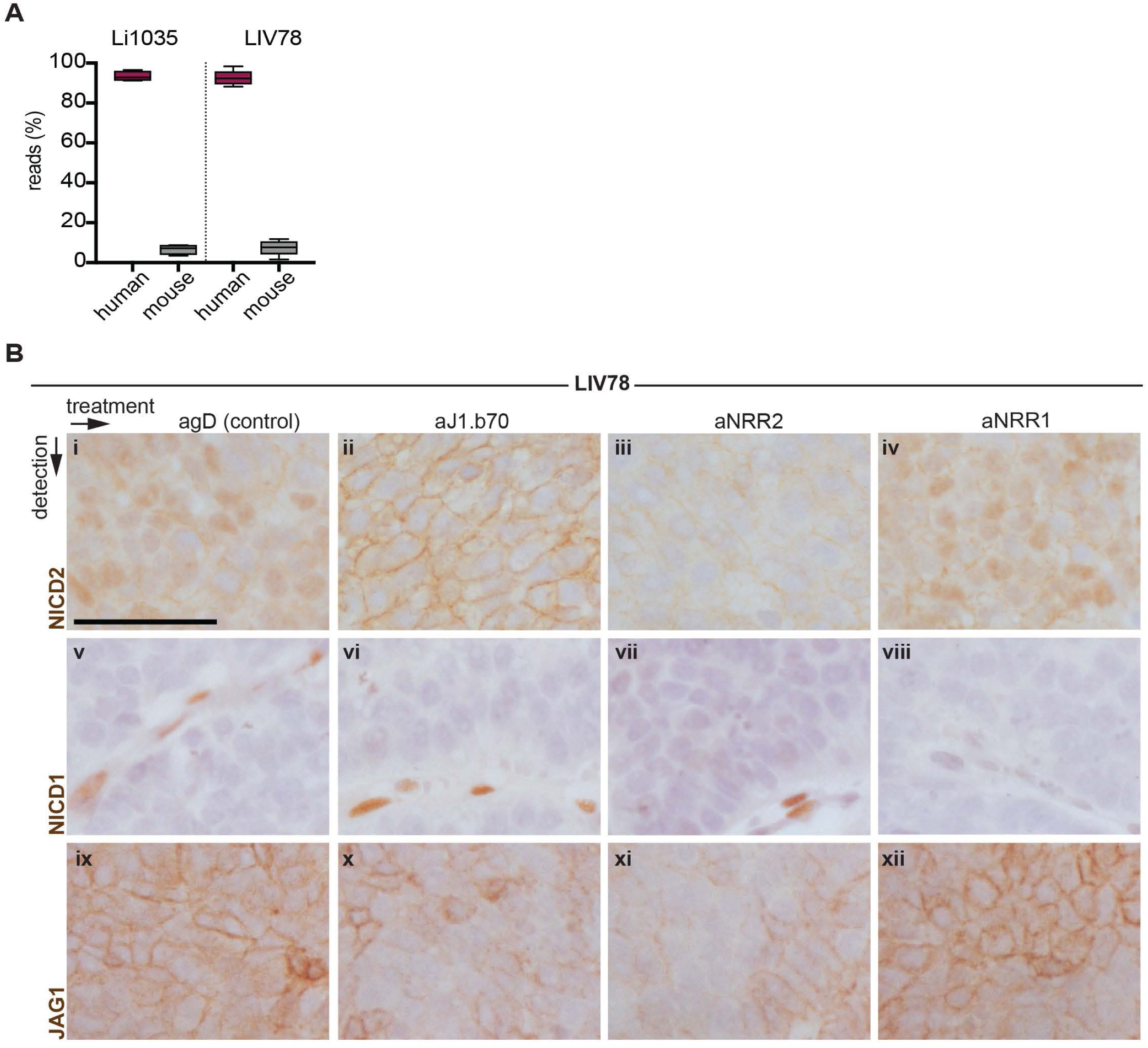
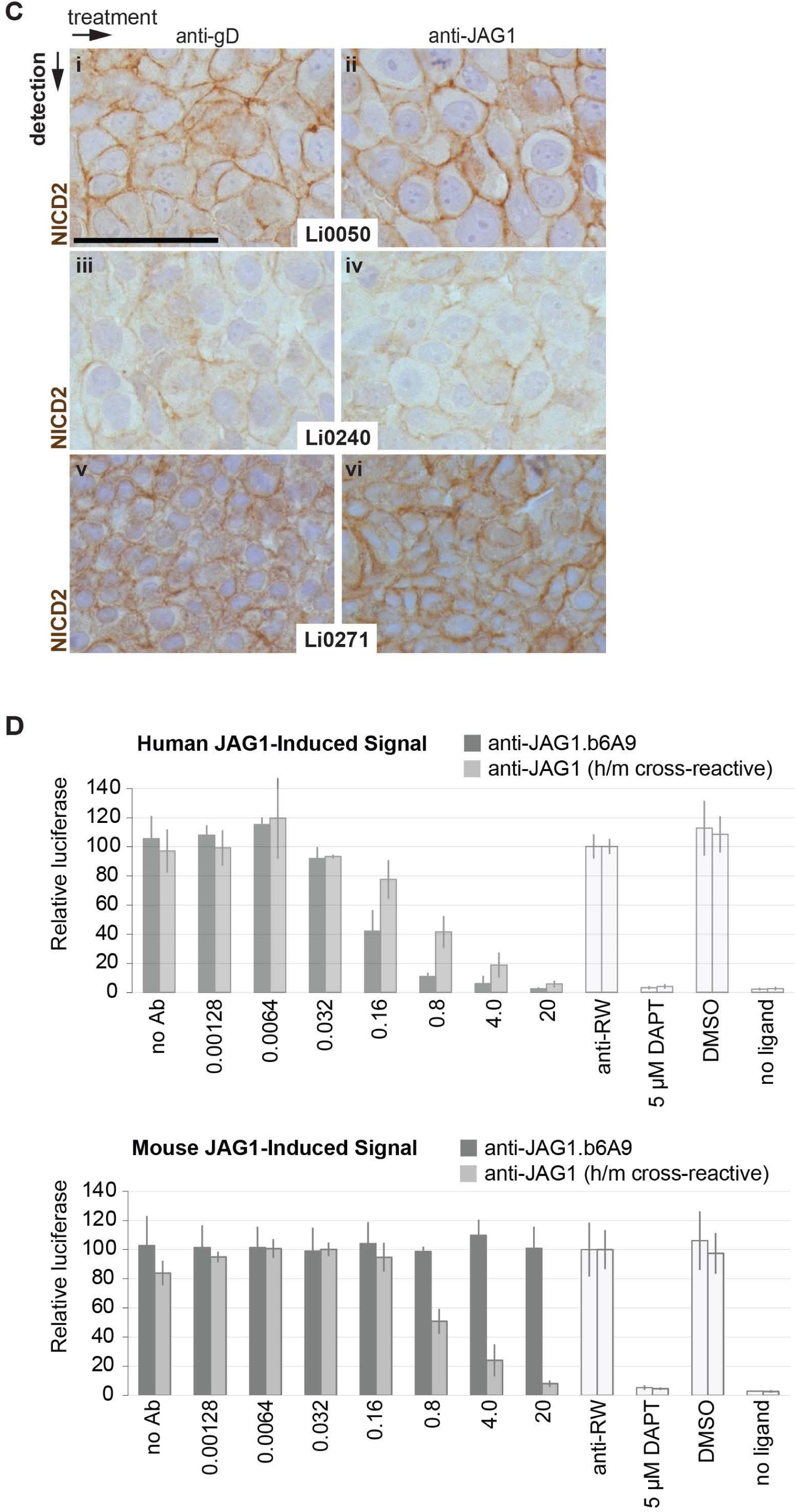
**A)** Percentages of human and mouse reads in PDX tumors used for bulk RNA sequencing. Data is shown for control antibody treated Li1035 (n=4) or LIV78 tumors (n=5). **B)** Immunohistochemical detection of NICD1, NRR2 and JAG1 in PDX model LIV78 treated with control antibodies or antibodies blocking JAG1 or the indicated receptor. Scale bar: 100µm **C)** Immunohistochemical detection of NRR2 in treatment-insensitive PDX models treated with control antibodies or antibodies blocking JAG1. Scale bar: 100µm **D)** Reporter assay showing selective binding of human but not mouse JAG1-induced signaling using aJ1.b6A9.

**Supplemental Figure 2 (goes with Main Figure3).**
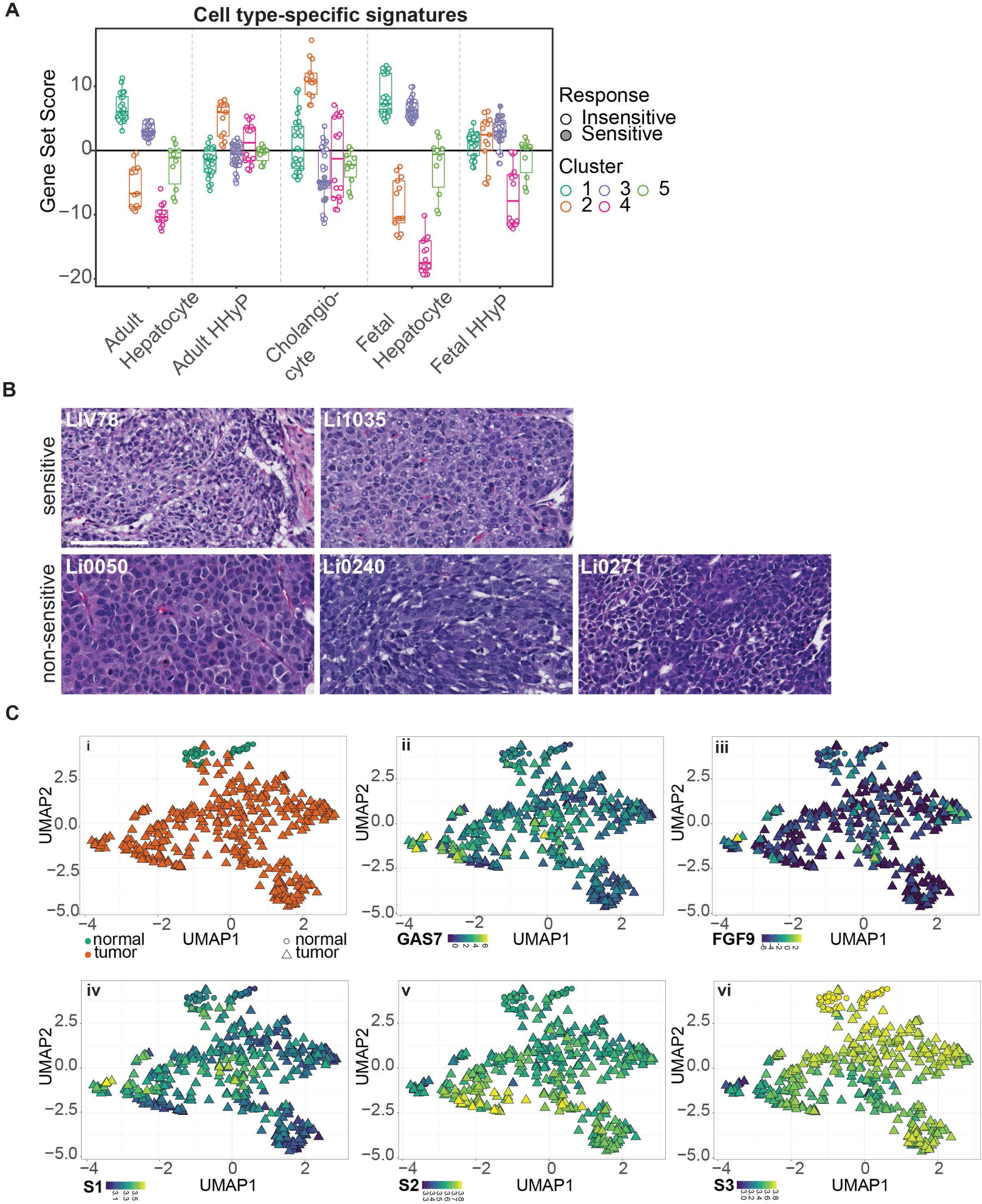
**A)** Expression of cell type specific signatures (Segal et al., 2019) in sensitive and non-sensitive models summarized by k-means cluster. **B)** Histological evaluation of JAG1-inhibition sensitive (top row) or non-sensitive (bottom row) tumors from hematoxylin and eosin stained sections of PDX tumors. All tumors are moderately-poorly differentiated HCCs with variable nuclear grade. Scale bar: 100µm. **C)** Two-dimensional representation of TCGA liver cancer/normal transcriptomes using UMAP projection. Each sample is represented by one data point and colored by sample type (i), Hoshida HCC subtype (ii - iv), and GAS7 (v) or FGF9 (vi) expression.

**Supplemental Figure 3 (goes with Main Figure4).**
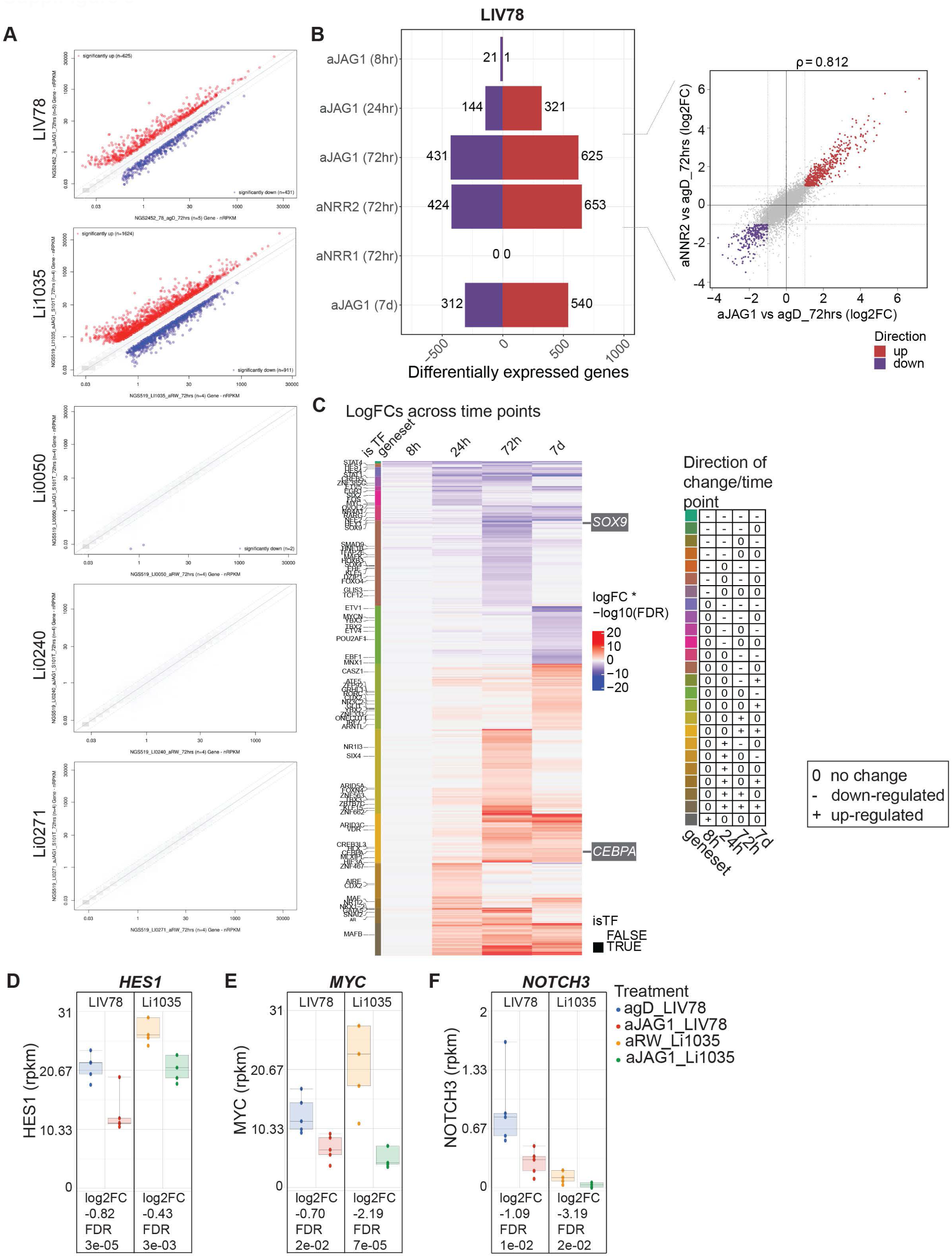
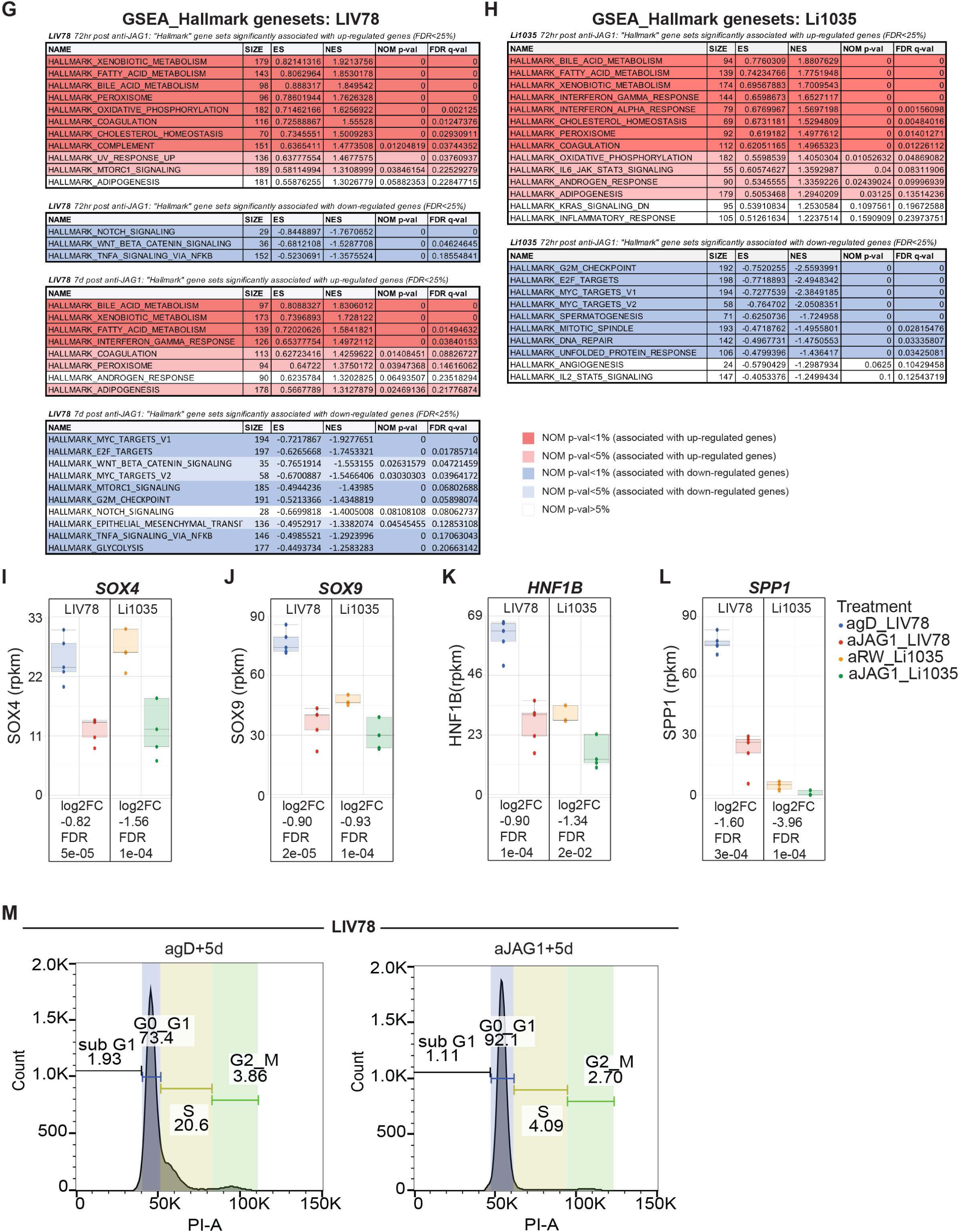
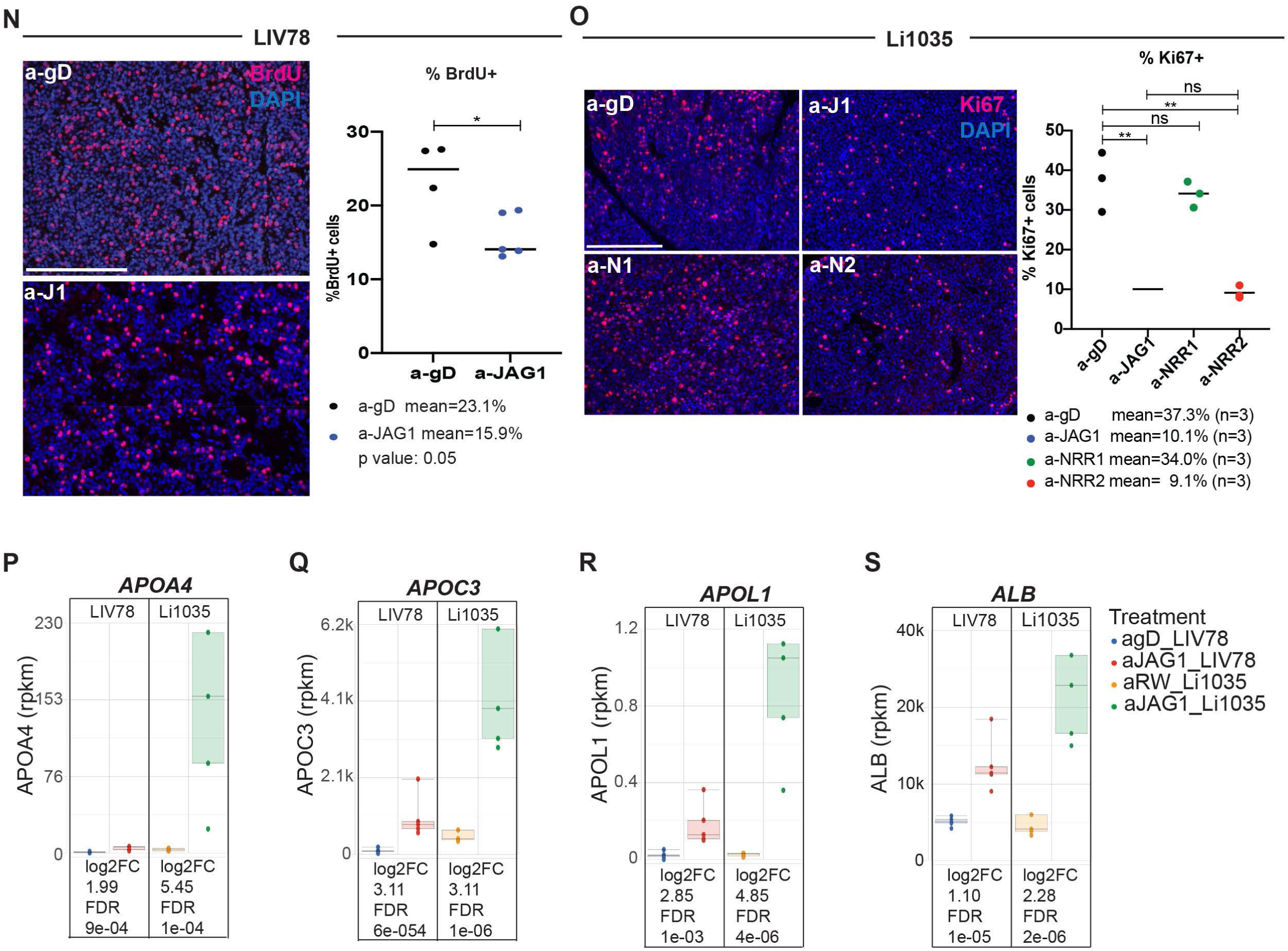
**A)** Scatterplots of differential gene expression in anti-J1.b70 treated tumors compared to control antibody treatment. Transcriptional responses 72 hours following single dose treatment are shown for regressing (top two panels) and non-regressing (bottom three panels) HCC PDX models. **B)** Summary of significant gene expression changes in LIV78 over time following treatment with anti-J1.b70. For the 72 hours post treatment time-point changes following NRR1 or NRR2 inhibition are shown for comparison. In (A) and (B) significantly changed transcripts [log_2_ fold change ≥|1|; adj.p-value >0.05] post single dose treatment are plotted. **C**) Heatmap showing differential gene expression over time for significantly up-/down-regulated genes. Transcription factors are highlighted on the left hand side and a color key provides the pattern of change across time points, which is summarized in the panel on the right. **D-F)** Expression of NOTCH target genes 72 hours after JAG1 inhibition in models LIV78 and Li1035 determined by bulk RNA sequencing. **G)** GSEA of differential expression between aJ1.b70 and control antibody treated LIV78 tumors at 72 hours and 7 days post treatment. **H)** GSEA of differential expression between aJ1.b70 and control antibody treated Li1035 tumors 72 hours after treatment. MSigDB hallmark gene sets were used for analyzes shown in (A) and (B). ES: enrichment score, NE: normalized enrichment score, FDR–false discovery rate. **I-L)** Expression of markers specific to cholangiocytes and bi-potent liver progenitors. **M)** Cell cycle analysis of LIV78 tumors 5 days following aJ1.b70 or control antibody treatment using propidium iodine staining and flow cytometry. Cycle histograms are shown for a representative sample for each group. In addition to confirming a treatment-induced change in cell cycle progression, this approach also allowed establishing baseline ploidy, which differs across HCC tumors and may be used as prognostic marker (Bou-Nader et al., 2020), LIV78 tumor cells were homogeneously diploid at baseline, and JAG1 inhibition did not result in a ploidy change 5 days post treatment. **N)** Quantification of Ki67 immunofluorescence staining in tissue sections of Li1035 tumors 3 days following treatment with aJ1b70, aNRR1, aNRR2, or control antibodies. Average percentage of Ki67+ cells from 3 frames (20x) per tumor are shown (n=3 per group). Scale bar: 100µm. **O)** Quantification of BrdU staining in tissue sections of LIV78 tumors 7 days following treatment with aJ1b70, or control antibody. BrdU was administered 1.5h prior to tumor harvest. Representative images are shown for each group (panels on left; scale bar: 100µm). Average percentage of BrdU+ cells from 3 frames (20x) per tumor are shown (panel on right; n=4 for agD,(control), n=5 for aJ1.b70).**P-R)** Expression of genes involved in fatty acid metabolism. **S)** Expression of the hepatocyte marker ALB. Data shown in I-L, and P-S was obtained by bulk RNA sequencing 72 hours after JAG1 inhibition in models LIV78 and Li1035.

**Supplemental Figure 4 (goes with Main Figure 5).**
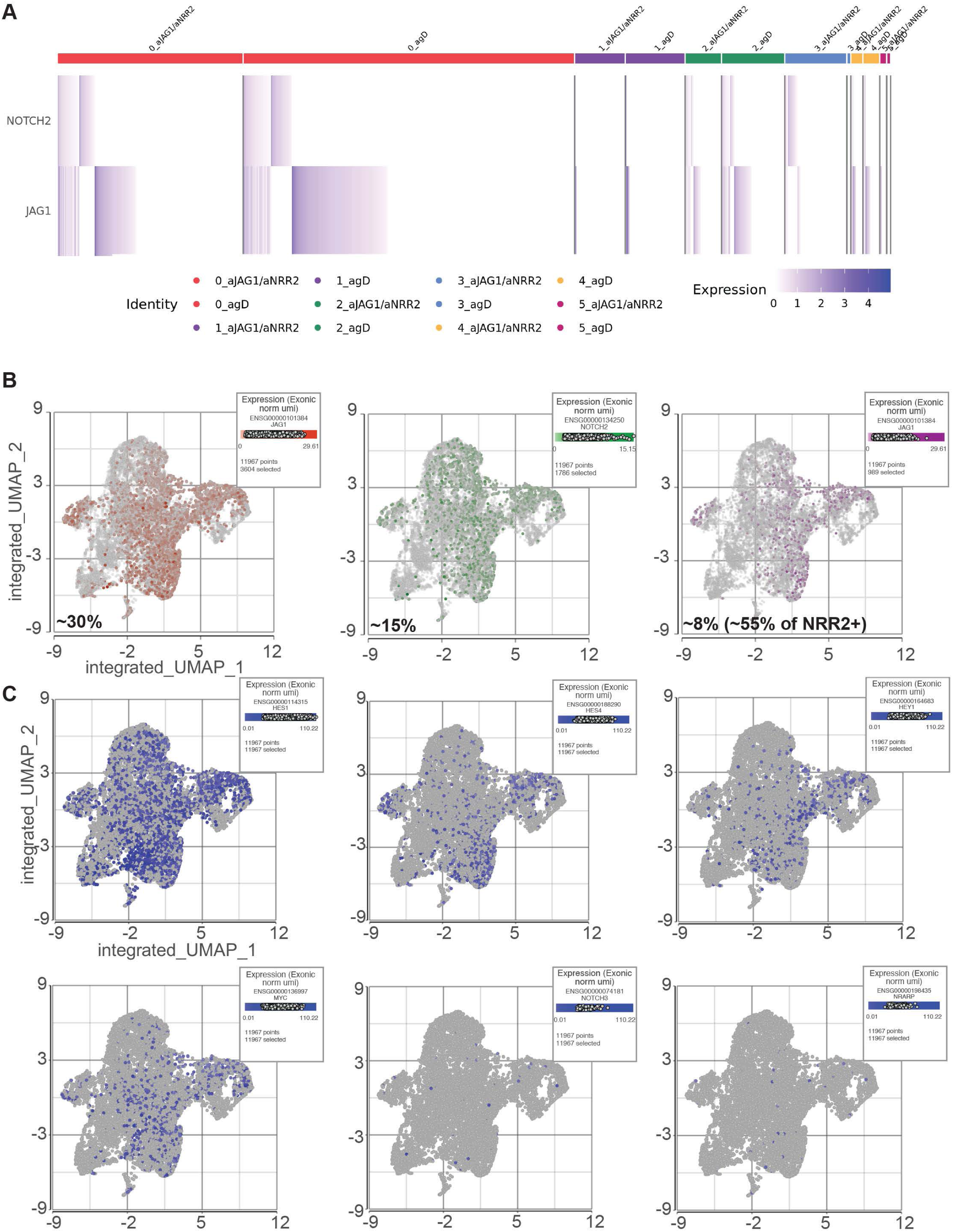
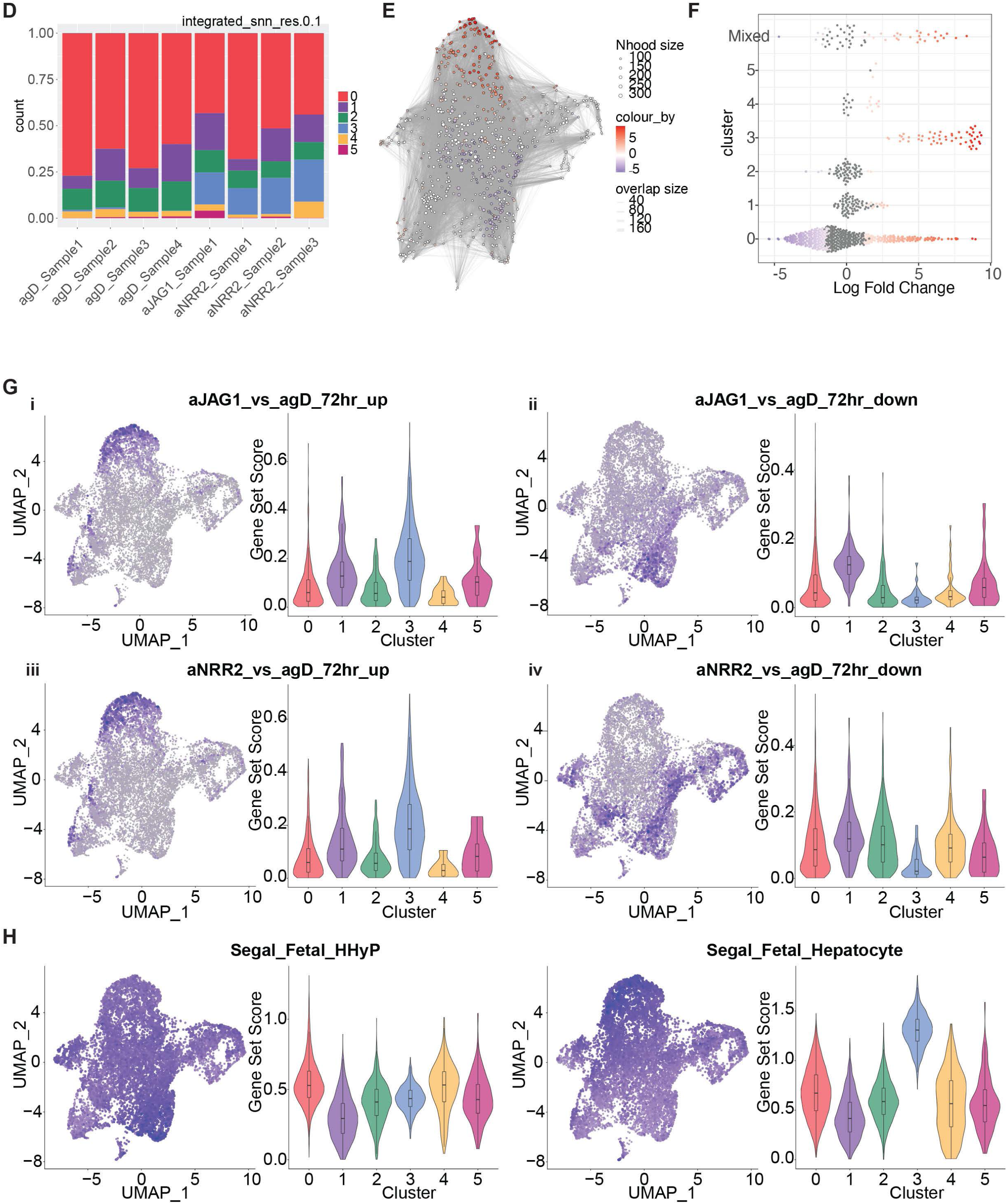
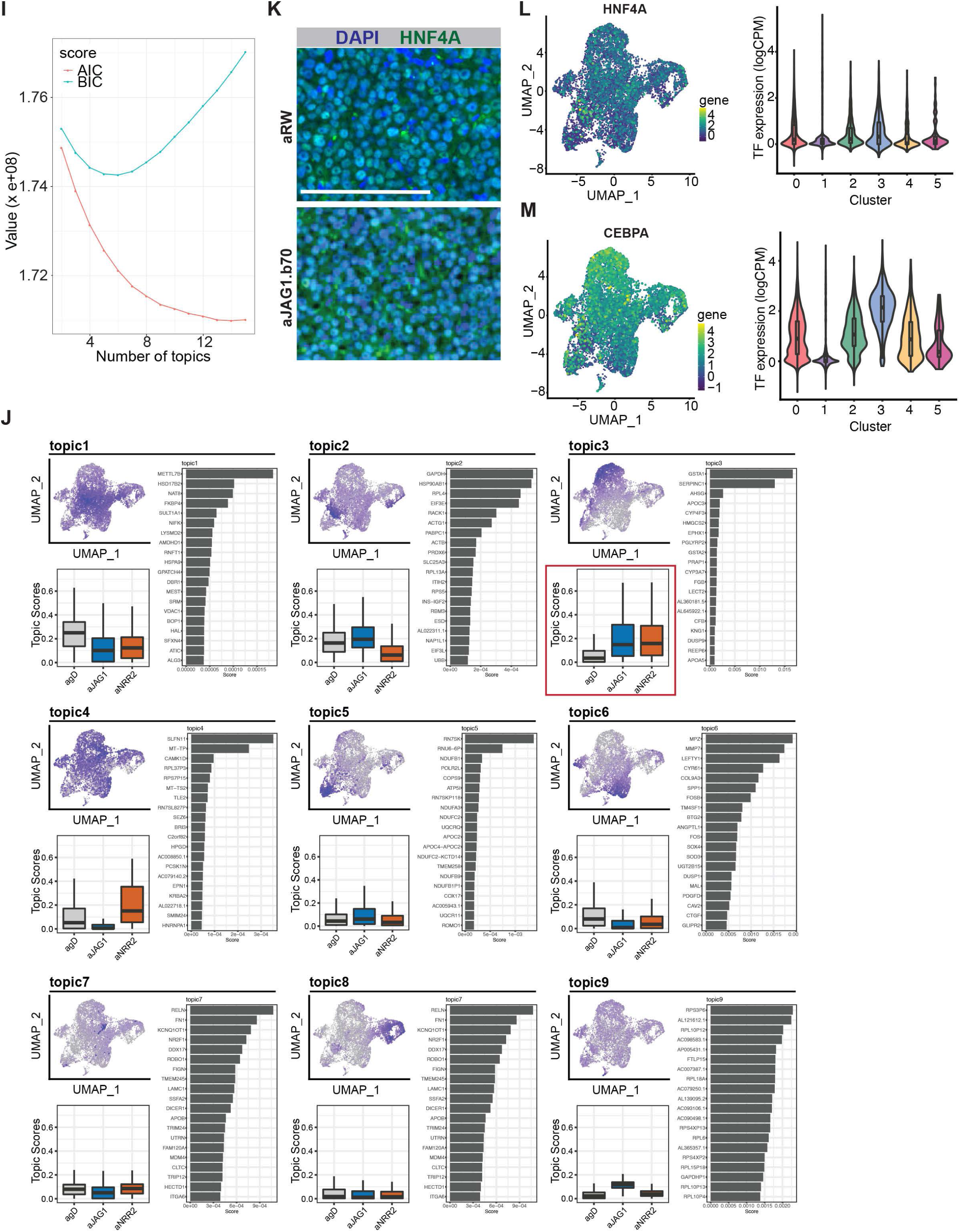
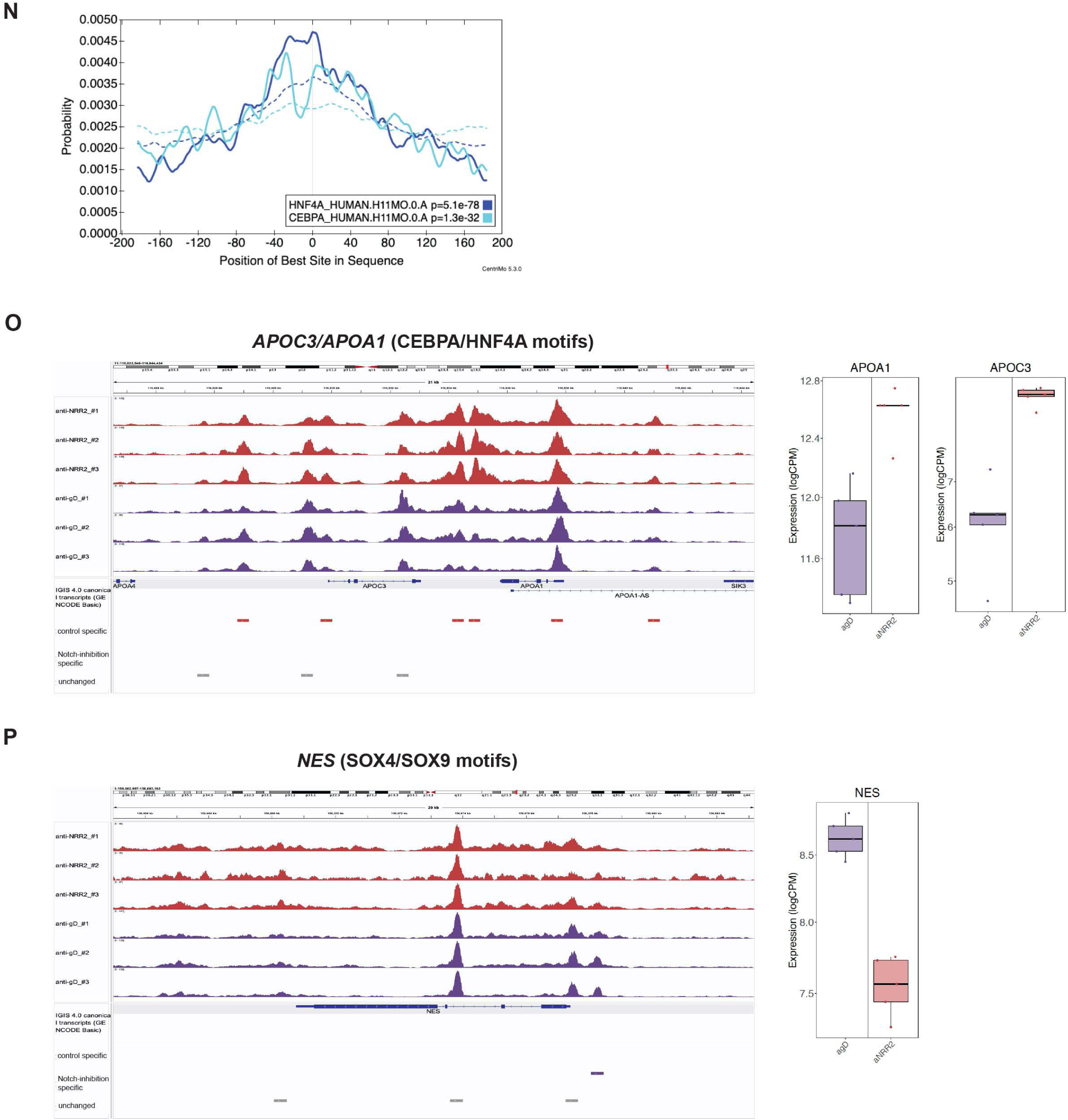
**A)** Expression of JAG1 and NOTCH2 in single cells arranged by UMAP cluster contribution and pathway inhibition status. aJ1.b70 and aNRR2-treated samples are summarized in one treatment group (aJAG1/aNRR2). **B)** UMAP plots of LIV78 tumor cell populations with JAG1 expressing cells (red) or NOTCH2 expressing cells (green) or cells expressing both JAG1 and NOTCH2 (purple) indicated. **C)** UMAP plots of LIV78 tumor cell populations with expression of common Notch pathway target genes indicated (blue). **D)** Neighborhood graph showing Milo differential abundance analysis between NOTCH inhibited and control samples. Each dot represents a neighborhood and is colored by its log fold change between conditions; circle sizes indicate the size of individual neighborhoods; graph edges represent the number of cells shared between neighborhoods. **E)** Beeswarm plot of the distribution of log fold change across clusters from figure 5A. Each dot shows one neighborhood. “Mixed” classification includes those neighborhoods belonging to multiple clusters. Colors represent the log fold change in abundance between NOTCH inhibited and control samples; neighborhoods with false discovery rate (FDR) > 0.1 are colored in gray. **F)** Proportion of cells contributing to individual UMAP clusters displayed for all individual samples **G)** Single cell expression of bulk RNA sequencing-derived signatures of significantly up-regulated (i,iii) or down-regulated (ii,iv) genes 3 days past JAG1(i,ii) or NRR2 (iii,iv) inhibition in LIV78 tumors. Single cell expression of each signature is shown in UMAP plots (left panels) and summarized for each cluster in violin plots (right panels). **H)** Single cell expression of cell type-specific signatures from *Segal et al* (Segal et al., 2019) in LIV78 tumors following Notch inhibition or control antibody treatment. UMAP plots and expression summaries for each single cell cluster as violin plots are shown for expression of a human fetal progenitor (left) and human fetal hepatocyte signature (right). **I)** AIC and BIC scores across a range [2-15] of topics. *K*=9 topics were determined to be the most informative. **J)** Topic modeling of LIV78 single cell data. For each topic a UMAP plot color-coded by the topic score (top left), a comparison of the topic score between treatment groups (bottom left) and the top 20 genes associated with the topic (right) are shown. **K)** Immunofluorescence detection of HNF4A expression on tissue sections from aJAG1.b70 or control antibody-treated Li1035 tumors (3days). Scale bar: 50µm. **L-M)** Single cell expression of *HNF4A* (L) or *CEBPA* (M) is shown in UMAP plot (left panels) and summarized for each cluster in violinplots (right panels). **N)** Centrimo analysis showing the location-specific preferential enrichment of HNF4A and CEBPA motifs. HNF4A binding shows centrally located enrichment, while CEBPA binding motifs show lateral enrichment in chromatin regions that are opening following NRR2 inhibition. **O)** Genome tracks of ATAC-sequencing signals in anti-NRR2-treated and control antibody treated LIV78 tumors across HNF4A- and CEBPA-bound promotors of genes associated with hepatocyte function (left panel) and their corresponding RNA-seq expression levels (right panel). **P)** Genome tracks of ATAC-sequencing signals in anti-NRR2-treated and control antibody treated LIV78 tumors in the SOX-transcription factor-bound promoter region of the progenitor marker NES (left panel) and corresponding RNA-seq-expression levels (right panel).

## Bibliography

Adams, J. M., & Jafar-Nejad, H. (2019). biomolecules The Roles of Notch Signaling in Liver Development and Disease. Biomolecules. 10.3390/biom9100608

Agnusdei, V., Minuzzo, S., Frasson, C., Grassi, A., Axelrod, F., Satyal, S., Gurney, A., Hoey, T., Seganfreddo, E., Basso, G., Valtorta, S., Moresco, R. M., Amadori, A., & Indraccolo, S. (2014). Therapeutic antibody targeting of Notch1 in T-acute lymphoblastic leukemia xenografts. Leukemia, 28(2), 278–288. 10.1038/LEU.2013.183

Aibar, S., González-Blas, C. B., Moerman, T., Huynh-Thu, V. A., Imrichova, H., Hulselmans, G., Rambow, F., Marine, J. C., Geurts, P., Aerts, J., Van Den Oord, J., Atak, Z. K., Wouters, J., & Aerts, S. (2017). SCENIC: single-cell regulatory network inference and clustering. Nature Methods, 14(11), 1083–1086. 10.1038/NMETH.4463

Akai, Y., Oitate, T., Koike, T., & Shiojiri, N. (2014). Impaired hepatocyte maturation, abnormal expression of biliary transcription factors and liver fibrosis in C/EBPα (Cebpa)-knockout mice. Histology and Histopathology, 29(1), 107–125. 10.14670/HH-29.107

Allen, F., & Maillard, I. (2021). Therapeutic Targeting of Notch Signaling: From Cancer to Inflammatory Disorders. Frontiers in Cell and Developmental Biology, 9. 10.3389/FCELL.2021.649205

Ally, A., Balasundaram, M., Carlsen, R., Chuah, E., Clarke, A., Dhalla, N., Holt, R. A., Jones, S. J. M., Lee, D., Ma, Y., Marra, M. A., Mayo, M., Moore, R. A., Mungall, A. J., Schein, J. E., Sipahimalani, P., Tam, A., Thiessen, N., Cheung, D., … Laird, P. W. (2017). Comprehensive and Integrative Genomic Characterization of Hepatocellular Carcinoma. Cell, 169(7), 1327–1341.e23. 10.1016/J.CELL.2017.05.046

Artavanis-Tsakonas, S., Rand, M. D., & Lake, R. J. (1999). Notch Signaling: Cell Fate Control and Signal Integration in Development. Science. https://www.science.org

Aster, J. C., Pear, W. S., & Blacklow, S. C. (2017). The Varied Roles of Notch in Cancer. Annual Review of Pathology, 12, 245–275. 10.1146/ANNUREV-PATHOL-052016-100127

Bachmann, K. A. (1996). The Cytochrome P450 Enzymes of Hepatic Drug Metabolism: How are their Activities Assessed In Vivo, and what is their Clinical Relevance? American Journal of Therapeutics, 3(2), 150–171. 10.1097/00045391-199602000-00009

Bailey, T. L., & MacHanick, P. (2012). Inferring direct DNA binding from ChIP-seq. Nucleic Acids Research, 40(17). 10.1093/NAR/GKS433

Bou-Nader, M., Caruso, S., Donne, R., Celton-Morizur, S., Calderaro, J., Gentric, G., Cadoux, M., L’Hermitte, A., Klein, C., Guilbert, T., Albuquerque, M., Couchy, G., Paradis, V., Couty, J. P., Zucman-Rossi, J., & Desdouets, C. (2020). Polyploidy spectrum: A new marker in HCC classification. Gut. 10.1136/gutjnl-2018-318021

Boyault, S., Rickman, D. S., De Reyniès, A., Balabaud, C., Rebouissou, S., Jeannot, E., Hérault, A., Saric, J., Belghiti, J., Franco, D., Bioulac-Sage, P., Laurent-Puig, P., & Zucman-Rossi, J. (2007). Transcriptome classification of HCC is related to gene alterations and to new therapeutic targets. Hepatology, 45(1), 42–52. 10.1002/HEP.21467

Calderaro, J., Ziol, M., Paradis, V., & Zucman-Rossi, J. (2019). Molecular and histological correlations in liver cancer. Journal of Hepatology, 71(3), 616–630. 10.1016/J.JHEP.2019.06.001

Choy, L., Hagenbeek, T. J., Solon, M., French, D., Finkle, D., Shelton, A., Venook, R., Brauer, M. J., & Siebel, C. W. (2017). Constitutive NOTCH3 Signaling Promotes the Growth of Basal Breast Cancers. Cancer Research, 77(6), 1439–1452. 10.1158/0008-5472.CAN-16-1022

Coffinier, C., Gresh, L., Fiette, L., Tronche, F., Schütz, G., Babinet, C., Pontoglio, M., Yaniv, M., & Barra, J. (2002). Bile system morphogenesis defects and liver dysfunction upon targeted deletion of HNF1beta. *Development (Cambridge*, England), 129(8), 1829–1838. 10.1242/DEV.129.8.1829

da Fonseca, L. G., Reig, M., & Bruix, J. (2020). Tyrosine Kinase Inhibitors and Hepatocellular Carcinoma. Clinics in Liver Disease, 24(4), 719–737. 10.1016/J.CLD.2020.07.012

Dann, E., Henderson, N. C., Teichmann, S. A., Morgan, M. D., & Marioni, J. C. (2022). Differential abundance testing on single-cell data using k-nearest neighbor graphs. Nature Biotechnology, 40(2), 245–253. 10.1038/S41587-021-01033-Z

De Obaldia, M. E., Bell, J. J., Wang, X., Harly, C., Yashiro-Ohtani, Y., Delong, J. H., Zlotoff, D. A., Sultana, D. A., Pear, W. S., & Bhandoola, A. (2013). T cell development requires constraint of the myeloid regulator C/EBP-α by the Notch target and transcriptional repressor Hes1. Nature Immunology, 14(12), 1277–1284. 10.1038/ni.2760

De Strooper, B., Annaert, W., Cupers, P., Saftig, P., Craessaerts, K., Mumm, J. S., Schroeter, E. H., Schrijvers, V., Wolfe, M. S., Ray, W. J., Goate, A., & Kopan, R. (1999). A presenilin-1-dependent gamma-secretase-like protease mediates release of Notch intracellular domain. Nature, 398(6727), 518–522. 10.1038/19083

Dey, K. K., Hsiao, C. J., & Stephens, M. (2017). Visualizing the structure of RNA-seq expression data using grade of membership models. PLoS Genetics, 13(3). 10.1371/JOURNAL.PGEN.1006599

Dill, M. T., Tornillo, L., Fritzius, T., Terracciano, L., Semela, D., Bettler, B., Heim, M. H., & Tchorz, J. S. (2013). Constitutive Notch2 signaling induces hepatic tumors in mice. *Hepatology (Baltimore*, Md.), 57(4), 1607–1619. 10.1002/HEP.26165

Fan, B., Malato, Y., Calvisi, D. F., Naqvi, S., Razumilava, N., Ribback, S., Gores, G. J., Dombrowski, F., Evert, M., Chen, X., & Willenbring, H. (2012). Cholangiocarcinomas can originate from hepatocytes in mice. Journal of Clinical Investigation, 122(8), 2911–2915. 10.1172/JCI63212

Ferrarotto, R., Mitani, Y., Diao, L., Guijarro, I., Wang, J., Zweidler-McKay, P., Bell, D., William, W. N., Glisson, B. S., Wick, M. J., Kapoun, A. M., Patnaik, A., Eckhardt, G., Munster, P., Faoro, L., Dupont, J., Lee, J. J., Futreal, A., El-Naggar, A. K., & Heymach, J. V. (2017). Activating NOTCH1 Mutations Define a Distinct Subgroup of Patients With Adenoid Cystic Carcinoma Who Have Poor Prognosis, Propensity to Bone and Liver Metastasis, and Potential Responsiveness to Notch1 Inhibitors. Journal of Clinical Oncology : Official Journal of the American Society of Clinical Oncology, 35(3), 352–360. 10.1200/JCO.2016.67.5264

Finn, R. S., Qin, S., Ikeda, M., Galle, P. R., Ducreux, M., Kim, T.-Y., Kudo, M., Breder, V., Merle, P., Kaseb, A. O., Li, D., Verret, W., Xu, D.-Z., Hernandez, S., Liu, J., Huang, C., Mulla, S., Wang, Y., Lim, H. Y., … Cheng, A.-L. (2020). Atezolizumab plus Bevacizumab in Unresectable Hepatocellular Carcinoma. The New England Journal of Medicine, 382(20), 1894–1905. 10.1056/NEJMOA1915745

Flodby, P., Barlow, C., Kylefjord, H., Ährlund-Richter, L., & Xanthopoulos, K. G. (1996). Increased hepatic cell proliferation and lung abnormalities in mice deficient in CCAAT/enhancer binding protein α. Journal of Biological Chemistry, 271(40), 24753–24760. 10.1074/jbc.271.40.24753

Forrest, W. F., Alicke, B., Mayba, O., Osinska, M., Jakubczak, M., Piatkowski, P., Choniawko, L., Starr, A., & Gould, S. E. (2020). Generalized additive mixed modeling of longitudinal tumor growth reduces bias and improves decision making in translational oncology. Volume 80*, Issue* 22, Pages 5089 - 5097, 80(22), 5089–5097. 10.1158/0008-5472.CAN-20-0342

Garcia-Alonso, L., Holland, C. H., Ibrahim, M. M., Turei, D., & Saez-Rodriguez, J. (2019). Benchmark and integration of resources for the estimation of human transcription factor activities. Genome Research, 29(8), 1363–1375. 10.1101/GR.240663.118

Goossens, N., Sun, X., & Hoshida, Y. (2015a). Molecular classification of hepatocellular carcinoma: potential therapeutic implications. Hepatic Oncology, 2(4), 371–379. 10.2217/hep.15.26

Goossens, N., Sun, X., & Hoshida, Y. (2015b). Molecular classification of hepatocellular carcinoma: potential therapeutic implications. Hepat. Oncol, 2(4), 371–379. 10.2217/hep.15.26

Gordon, W. R., Zimmerman, B., He, L., Miles, L. J., Huang, J., Tiyanont, K., McArthur, D. G., Aster, J. C., Perrimon, N., Loparo, J. J., & Blacklow, S. C. (2015). Mechanical Allostery: Evidence for a Force Requirement in the Proteolytic Activation of Notch. Developmental Cell, 33(6), 729–736. 10.1016/J.DEVCEL.2015.05.004

Holland, C. H., Szalai, B., & Saez-Rodriguez, J. (2020). Transfer of regulatory knowledge from human to mouse for functional genomics analysis. Biochimica et Biophysica Acta. Gene Regulatory Mechanisms, 1863(6). 10.1016/J.BBAGRM.2019.194431

Holland, C. H., Tanevski, J., Perales-Patón, J., Gleixner, J., Kumar, M. P., Mereu, E., Joughin, B. A., Stegle, O., Lauffenburger, D. A., Heyn, H., Szalai, B., & Saez-Rodriguez, J. (2020). Robustness and applicability of transcription factor and pathway analysis tools on single-cell RNA-seq data. Genome Biology, 21(1). 10.1186/S13059-020-1949-Z

Hoshida, Y., Nijman, S. M. B., Kobayashi, M., Chan, J. A., Brunet, J. P., Chiang, D. Y., Villanueva, A., Newell, P., Ikeda, K., Hashimoto, M., Watanabe, G., Gabriel, S., Friedman, S. L., Kumada, H., Llovet, J. M., & Golub, T. R. (2009). Integrative transcriptome analysis reveals common molecular subclasses of human hepatocellular carcinoma. Cancer Research. 10.1158/0008-5472.CAN-09-1089

Huntzicker, E. G., Hötzel, K., Choy, L., Che, L., Ross, J., Pau, G., Sharma, N., Siebel, C. W., Chen, X., & French, D. M. (2015a). Differential effects of targeting Notch receptors in a mouse model of liver cancer. Hepatology. 10.1002/hep.27566

Huntzicker, E. G., Hötzel, K., Choy, L., Che, L., Ross, J., Pau, G., Sharma, N., Siebel, C. W., Chen, X., & French, D. M. (2015b). Differential effects of targeting Notch receptors in a mouse model of liver cancer. Hepatology, 61(3), 942–952. 10.1002/hep.27566

Kawaguchi, K., Honda, M., Yamashita, T., Okada, H., Shirasaki, T., Nishikawa, M., Nio, K., Arai, K., Sakai, Y., Yamashita, T., Mizukoshi, E., & Kaneko, S. (2016). Jagged1 DNA Copy Number Variation Is Associated with Poor Outcome in Liver Cancer. The American Journal of Pathology, 186(8), 2055–2067. 10.1016/J.AJPATH.2016.04.011

Klinakis, A., Lobry, C., Abdel-Wahab, O., Oh, P., Haeno, H., Buonamici, S., Van De Walle, I., Cathelin, S., Trimarchi, T., Araldi, E., Liu, C., Ibrahim, S., Beran, M., Zavadil, J., Efstratiadis, A., Taghon, T., Michor, F., Levine, R. L., & Aifantis, I. (2011). A novel tumour-suppressor function for the Notch pathway in myeloid leukaemia. Nature, 473(7346), 230–233. 10.1038/nature09999

Koch, U., Lehal, R., & Radtke, F. (2013). Stem cells living with a Notch. *Development (Cambridge*, England), 140(4), 689–704. 10.1242/DEV.080614

Kulakovskiy, I. V., Vorontsov, I. E., Yevshin, I. S., Sharipov, R. N., Fedorova, A. D., Rumynskiy, E. I., Medvedeva, Y. A., Magana-Mora, A., Bajic, V. B., Papatsenko, D. A., Kolpakov, F. A., & Makeev, V. J. (2018). HOCOMOCO: towards a complete collection of transcription factor binding models for human and mouse via large-scale ChIP-Seq analysis. Nucleic Acids Research, 46(D1), D252–D259. 10.1093/NAR/GKX1106

Lafkas, D., Shelton, A., Chiu, C., de Leon boenig, G., Chen, Y., Stawicki, S. S., Siltanen, C., Reichelt, M., Zhou, M., Wu, X., Eastham-Anderson, J., Moore, H., Roose-Girma, M., Chinn, Y., Hang, J. Q., Warming, S., Egen, J., Lee, W. P., Austin, C., … Siebel, christian W. (2015). Therapeutic antibodies reveal Notch control of transdifferentiation in the adult lung. Nature. 10.1038/nature15715

Law, C. W., Chen, Y., Shi, W., & Smyth, G. K. (2014). voom: Precision weights unlock linear model analysis tools for RNA-seq read counts. Genome Biology, 15(2). 10.1186/GB-2014-15-2-R29

Lesaffer, B., Verboven, E., Huffel, L. Van, Moya, I. M., Grunsven, L. A. van, Leclercq, I. A., Lemaigre, F. P., & Halder, G. (2019). Comparison of the Opn-CreER and Ck19-CreER Drivers in Bile Ducts of Normal and Injured Mouse Livers. Cells, 8(4), 380. 10.3390/CELLS8040380

Liberzon, A., Birger, C., Thorvaldsdóttir, H., Ghandi, M., Mesirov, J. P., & Tamayo, P. (2015). The Molecular Signatures Database (MSigDB) hallmark gene set collection. Cell Systems, 1(6), 417. 10.1016/J.CELS.2015.12.004

Liberzon, A., Subramanian, A., Pinchback, R., Thorvaldsdóttir, H., Tamayo, P., & Mesirov, J. P. (2011). Molecular signatures database (MSigDB) 3.0. *Bioinformatics (Oxford*, England), 27(12), 1739–1740. 10.1093/BIOINFORMATICS/BTR260

Lim, J. S., Ibaseta, A., Fischer, M. M., Cancilla, B., O’Young, G., Cristea, S., Luca, V. C., Yang, Di., Jahchan, N. S., Hamard, C., Antoine, M., Wislez, M., Kong, C., Cain, J., Liu, Y. W., Kapoun, A. M., Garcia, K. C., Hoey, T., Murriel, C. L., & Sage, J. (2017). Intratumoural heterogeneity generated by Notch signalling promotes small-cell lung cancer. Nature. 10.1038/nature22323

Llovet, J. M., Castet, F., Heikenwalder, M., Maini, M. K., Mazzaferro, V., Pinato, D. J., Pikarsky, E., Zhu, A. X., & Finn, R. S. (2022). Immunotherapies for hepatocellular carcinoma. Nature Reviews Clinical Onocology. 10.1038/s41571-021-00573-2

Long, J. E., Wongchenko, M. J., Nickles, D., Chung, W. J., Wang, B. er, Riegler, J., Li, J., Li, Q., Sandoval, W., Eastham-Anderson, J., Modrusan, Z., Junttila, T., Carano, R. A. D., Foreman, O., Yan, Y., & Junttila, M. R. (2019). Therapeutic resistance and susceptibility is shaped by cooperative multi-compartment tumor adaptation. Cell Death and Differentiation, 26(11), 2416–2429. 10.1038/S41418-019-0310-0

Loomes, K. M., Underkoffler, L. A., Morabito, J., Gottlieb, S., Piccoli, D. A., Spinner, N. B., Baldwin, H. S., & Oakey, R. J. (1999). The expression of Jagged1 in the developing mammalian heart correlates with cardiovascular disease in Alagille syndrome. Human Molecular Genetics, 8(13), 2443–2449. 10.1093/HMG/8.13.2443

Mack, J. J., & Luisa Iruela-Arispe, M. (2018). NOTCH regulation of the endothelial cell phenotype. Current Opinion in Hematology, 25(3), 212–218. 10.1097/MOH.0000000000000425

Marx, V. (2015). Cancer: A most exceptional response. Nature, 520(7547), 389–393. 10.1038/520389A

McCright, B., Lozier, J., & Gridley, T. (2002). A mouse model of Alagille syndrome: Notch2 as a genetic modifier of Jag1 haploinsufficiency. In Development (Vol. 129, Issue 4, pp. 1075–1082). Development. https://pubmed.ncbi.nlm.nih.gov/11861489/

McDavid, A., Finak, G., Chattopadyay, P. K., Dominguez, M., Lamoreaux, L., Ma, S. S., Roederer, M., & Gottardo, R. (2013). Data exploration, quality control and testing in single-cell qPCR-based gene expression experiments. *Bioinformatics (Oxford*, England), 29(4), 461–467. 10.1093/BIOINFORMATICS/BTS714

McLeay, R. C., & Bailey, T. L. (2010). Motif Enrichment Analysis: a unified framework and an evaluation on ChIP data. BMC Bioinformatics, 11. 10.1186/1471-2105-11-165

Mederacke, I., Dapito, D. H., Affò, S., Uchinami, H., & Schwabe, R. F. (2015). High-yield and high-purity isolation of hepatic stellate cells from normal and fibrotic mouse livers. Nature Protocols, 10(2), 305–315. 10.1038/NPROT.2015.017

Parviz, F., Matullo, C., Garrison, W. D., Savatski, L., Adamson, J. W., Ning, G., Kaestner, K. H., Rossi, J. M., Zaret, K. S., & Duncan, S. A. (2003). Hepatocyte nuclear factor 4α controls the development of a hepatic epithelium and liver morphogenesis. Nature Genetics, 34(3), 292–296. 10.1038/ng1175

Poncy, A., Antoniou, A., Cordi, S., Pierreux, C. E., Jacquemin, P., & Lemaigre, F. P. (2015). Transcription factors SOX4 and SOX9 cooperatively control development of bile ducts. Developmental Biology, 404(2), 136–148. 10.1016/J.YDBIO.2015.05.012

Prior, N., Hindley, C. J., Rost, F., Meléndez, E., Lau, W. W. Y., Göttgens, B., Rulands, S., Simons, B. D., & Huch, M. (2019). Lgr5+ stem and progenitor cells reside at the apex of a heterogeneous embryonic hepatoblast pool. Development (Cambridge). 10.1242/dev.174557

Ramsay, E. E., & Dilda, P. J. (2014). Glutathione S-conjugates as prodrugs to target drug-resistant tumors. Frontiers in Pharmacology, 5. 10.3389/FPHAR.2014.00181

Ritchie, M. E., Phipson, B., Wu, D., Hu, Y., Law, C. W., Shi, W., & Smyth, G. K. (2015). limma powers differential expression analyses for RNA-sequencing and microarray studies. Nucleic Acids Research, 43(7), e47. 10.1093/NAR/GKV007

Ross-Innes, C. S., Stark, R., Teschendorff, A. E., Holmes, K. A., Ali, H. R., Dunning, M. J., Brown, G. D., Gojis, O., Ellis, I. O., Green, A. R., Ali, S., Chin, S. F., Palmieri, C., Caldas, C., & Carroll, J. S. (2012). Differential oestrogen receptor binding is associated with clinical outcome in breast cancer. Nature, 481(7381), 389–393. 10.1038/NATURE10730

Segal, J. M., Kent, D., Wesche, D. J., Ng, S. S., Serra, M., Oulès, B., Kar, G., Emerton, G., Blackford, S. J. I., Darmanis, S., Miquel, R., Luong, T. V., Yamamoto, R., Bonham, A., Jassem, W., Heaton, N., Vigilante, A., King, A., Sancho, R., … Rashid, S. T. (2019). Single cell analysis of human foetal liver captures the transcriptional profile of hepatobiliary hybrid progenitors. Nature Communications. 10.1038/s41467-019-11266-x

Si-Tayeb, K., Lemaigre, F. P., & Duncan, S. A. (2010). Organogenesis and Development of the Liver. Developmental Cell, 18(2), 175–189. 10.1016/J.DEVCEL.2010.01.011

Siebel, C., & Lendahl, U. (2017). Notch Signaling in Development, Tissue Homeostasis, and Disease. Physiological Reviews, 97(4), 1235–1294. 10.1152/PHYSREV.00005.2017

Siolas, D., & Hannon, G. J. (2013). Patient-Derived Tumor Xenografts: Transforming Clinical Samples into Mouse Models. 10.1158/0008-5472.CAN-13-1069 Dann, E., Henderson, N. C., Teichmann, S. A., Morgan, M. D., & Marioni, J. C. (2022). Differential abundance testing on single-cell data using k-nearest neighbor graphs. Nature Biotechnology, 40(2), 245–253. 10.1038/S41587-021-01033-Z

Smith, D. C., Chugh, R., Patnaik, A., Papadopoulos, K. P., Wang, M., Kapoun, A. M., Xu, L., Dupont, J., Stagg, R. J., & Tolcher, A. (2019). A phase 1 dose escalation and expansion study of Tarextumab (OMP-59R5) in patients with solid tumors. Investigational New Drugs, 37(4), 722–730. 10.1007/S10637-018-0714-6

Srinivasan, K., Friedman, B. A., Larson, J. L., Lauffer, B. E., Goldstein, L. D., Appling, L. L., Borneo, J., Poon, C., Ho, T., Cai, F., Steiner, P., Van Der Brug, M. P., Modrusan, Z., Kaminker, J. S., & Hansen, D. V. (2016). Untangling the brain’s neuroinflammatory and neurodegenerative transcriptional responses. Nature Communications, 7. 10.1038/NCOMMS11295

Stanger, B. Z. (2015). Cellular homeostasis and repair in the mammalian liver. Annual Review of Physiology, 77, 179–200. 10.1146/ANNUREV-PHYSIOL-021113-170255

Stark, R. (University of C., & Brown, G. (University of C. (2011). DiffBind:Differential binding analysis of ChIP-Seq peak data. Bioconductor data.

Stefflova, K., Thybert, D., Wilson, M. D., Streeter, I., Aleksic, J., Karagianni, P., Brazma, A., Adams, D. J., Talianidis, I., Marioni, J. C., Flicek, P., & Odom, D. T. (2013). XCooperativity and rapid evolution of cobound transcription factors in closely related mammals. Cell, 154(3). 10.1016/j.cell.2013.07.007

Stoeck, A., Lejnine, S., Truong, A., Pan, L., Wang, H., Zang, C., Yuan, J., Ware, C., Maclean, J., Garrett-Engele, P. W., Kluk, M., Laskey, J., Haines, B. B., Moskaluk, C., Zawel, L., Fawell, S., Gilliland, G., Zhang, T., Kremer, B. E., … Sathyanarayanan, S. (2014). Discovery of Biomarkers Predictive of GSI Response in Triple-Negative Breast cancer and Adenoid Cystic carcinoma. Cancer Discovery. 10.1158/2159-8290.CD-13-0830

Stuart, T., Butler, A., Hoffman, P., Hafemeister, C., Papalexi, E., Mauck, W. M., Hao, Y., Stoeckius, M., Smibert, P., & Satija, R. (2019). Comprehensive Integration of Single-Cell Data. Cell, 177(7), 1888–1902.e21. 10.1016/J.CELL.2019.05.031

Subramanian, A., Tamayo, P., Mootha, V. K., Mukherjee, S., Ebert, B. L., Gillette, M. A., Paulovich, A., Pomeroy, S. L., Golub, T. R., Lander, E. S., & Mesirov, J. P. (2005). Gene set enrichment analysis: A knowledge-based approach for interpreting genome-wide expression profiles. Proceedings of the National Academy of Sciences of the United States of America, 102(43), 15545–15550. 10.1073/pnas.0506580102

Sung, H., Ferlay, J., Siegel, R., Laversanne, M., Soerjomataram, I., Jemal, A., & Bray, F. (2021). Global Cancer Statistics 2020: GLOBOCAN Estimates of Incidence and Mortality Worldwide for 36 Cancers in 185 Countries. 10.3322/caac.21660

Tachmatzidi, E. C., Galanopoulou, O., Talianidis, I., & Kalyuzhny, E. (2021). Transcription Control of Liver Development. 10.3390/cells10082026

Tanimizu, N., & Miyajima, A. (2004). Notch signaling controls hepatoblast differentiation by altering the expression of liver-enriched transcription factors. In Journal of Cell Science (Vol. 117, Issue 15, pp. 3165–3174). 10.1242/jcs.01169

Trefts, E., Gannon, M., & Wasserman, D. H. (2017). The liver. Current Biology, 27(21), R1147–R1151. 10.1016/J.CUB.2017.09.019

Viatour, P., Ehmer, U., Saddic, L. A., Dorrell, C., Andersen, J. B., Lin, C., Zmoos, A. F., Mazur, P. K., Schaffer, B. E., Ostermeier, A., Vogel, H., Sylvester, K. G., Thorgeirsson, S. S., Grompe, M., & Sage, J. (2011). Notch signaling inhibits hepatocellular carcinoma following inactivation of the RB pathway. The Journal of Experimental Medicine, 208(10), 1963–1976. 10.1084/JEM.20110198

Villanueva, A., Alsinet, C., Yanger, K., Hoshida, Y., Zong, Y., Toffanin, S., Rodriguez-Carunchio, L., Solé, M., Thung, S., Stanger, B. Z., & Llovet, J. M. (2012). Notch signaling is activated in human hepatocellular carcinoma and induces tumor formation in mice. Gastroenterology, 143(6). 10.1053/j.gastro.2012.09.002

Wang, N. D., Finegold, M. J., Bradley, A., Ou, C. N., Abdelsayed, S. V., Wilde, M. D., Taylor, L. R., Wilson, D. R., & Darlington, G. J. (1995). Impaired energy homeostasis in C/EBPα knockout mice. Science, 269(5227), 1108–1112. 10.1126/science.7652557

Watt, A. J., Garrison, W. D., & Duncan, S. A. (2003). HNF4: A central regulator of hepatocyte differentiation and function. In Hepatology (Vol. 37, Issue 6, pp. 1249– 1253). W.B. Saunders. 10.1053/jhep.2003.50273

Wheeler, D. A., Takebe, N., Hinoue, T., Hoadley, K. A., Cardenas, M. F., Hamilton, A. M., Laird, P. W., Wang, L., Johnson, A., Dewal, N., Miller, V., Piñeyro, D., Castro de Moura, M., Esteller, M., Shen, H., Zenklusen, J. C., Tarnuzzer, R., McShane, L. M., Tricoli, J. V., … Staudt, L. M. (2021). Molecular Features of Cancers Exhibiting Exceptional Responses to Treatment. Cancer Cell, 39(1), 38–53.e7. 10.1016/J.CCELL.2020.10.015

Wu, T. D., & Nacu, S. (2010). Fast and SNP-tolerant detection of complex variants and splicing in short reads. *Bioinformatics (Oxford*, England), 26(7), 873–881. 10.1093/BIOINFORMATICS/BTQ057

Wu, T., Hu, E., Xu, S., Chen, M., Guo, P., Dai, Z., Feng, T., Zhou, L., Tang, W., Zhan, L., Fu, X., Liu, S., Bo, X., & Yu, G. (2021). clusterProfiler 4.0: A universal enrichment tool for interpreting omics data. Innovation (Cambridge (Mass.)), 2(3). 10.1016/J.XINN.2021.100141

Wu, Y., Cain-Hom, C., Choy, L., Hagenbeek, T. J., De Leon, G. P., Chen, Y., Finkle, D., Venook, R., Wu, X., Ridgway, J., Schahin-Reed, D., Dow, G. J., Shelton, A., Stawicki, S., Watts, R. J., Zhang, J., Choy, R., Howard, P., Kadyk, L., … Siebel, C. W. (2010). Therapeutic antibody targeting of individual Notch receptors. Nature. 10.1038/nature08878

Xiao, Y., Hsiao, T. H., Suresh, U., Chen, H. I. H., Wu, X., Wolf, S. E., & Chen, Y. (2014). A novel significance score for gene selection and ranking. *Bioinformatics (Oxford*, England), 30(6), 801–807. 10.1093/BIOINFORMATICS/BTR671

Xu, H., & Wang, L. (2021). The Role of Notch Signaling Pathway in Non-Alcoholic Fatty Liver Disease. In Frontiers in Molecular Biosciences (Vol. 8). 10.3389/fmolb.2021.792667

Yamasaki, H., Sada, A., Iwata, T., Niwa, T., Tomizawa, M., Xanthopoulos, K. G., Koike, T., & Shiojiri, N. (2006). Suppression of C/EBPα expression in periportal hepatoblasts may stimulate biliary cell differentiation through increased Hnf6 and Hnf1b expression. Development, 133(21), 4233–4243. 10.1242/dev.02591

Yimlamai, D., Christodoulou, C., Galli, G. G., Yanger, K., Pepe-Mooney, B., Gurung, B., Shrestha, K., Cahan, P., Stanger, B. Z., & Camargo, F. D. (2014). Hippo pathway activity influences liver cell fate. Cell, 157(6). 10.1016/j.cell.2014.03.060

Zhu, C., Ho, Y. J., Salomao, M. A., Dapito, D. H., Bartolome, A., Schwabe, R. F., Lee, J. S., Lowe, S. W., & Pajvani, U. B. (2021). Notch activity characterizes a common hepatocellular carcinoma subtype with unique molecular and clinicopathologic features. Journal of Hepatology, 74(3), 613–626. 10.1016/J.JHEP.2020.09.032

Zhu, C., Kim, K. J., Wang, X., Bartolome, A., Salomao, M., Dongiovanni, P., Meroni, M., Graham, M. J., Yates, K. P., Diehl, A. M., Schwabe, R. F., Tabas, I., Valenti, L., Lavine, J. E., & Pajvani, U. B. (2018). Hepatocyte Notch activation induces liver fibrosis in nonalcoholic steatohepatitis. Science Translational Medicine, 10(468). 10.1126/SCITRANSLMED.AAT0344

Zong, Y., Panikkar, A., Xu, J., Antoniou, A., Raynaud, P., Lemaigre, F., & Stanger, B. Z. (2009). Notch signaling controls liver development by regulating biliary differentiation. Development, 136(10), 1727–1739. 10.1242/dev.029140

